# Thermodynamically Consistent Estimation of Gibbs Free Energy from Data: Data Reconciliation Approach

**DOI:** 10.1101/492819

**Authors:** Saman Salike, Nirav Bhatt

## Abstract

**Motivation:** Thermodynamic analysis of biological reaction networks requires the availability of accurate and consistent values of Gibbs free energies of reaction and formation. These Gibbs energies can be measured directly via the careful design of experiments or can be computed from the curated Gibbs free energy databases. However, the computed Gibbs free energies of reactions and formations do not satisfy the thermodynamic constraints due to the compounding effect of measurement errors in the experimental data. The propagation of these errors can lead to a false prediction of pathway feasibility and uncertainty in the estimation of thermodynamic parameters.

**Results:** This work proposes a data reconciliation framework for thermodynamically consistent estimation of Gibbs free energies of reaction, formation and group contributions from experimental data. In this framework, we formulate constrained optimization problems that reduce measurement errors and their effects on the estimation of Gibbs energies such that the thermodynamic constraints are satisfied. When a subset of Gibbs free energies of formations is unavailable, it is shown that the accuracy of their resulting estimates is better than that of existing empirical prediction methods. Moreover, we also show that the estimation of group contributions can be improved using this approach. Further, we provide guidelines based on this approach for performing systematic experiments to estimate unknown Gibbs formation energies.

**Availability:** The MATLAB code for the executing the proposed algorithm is available for free on the GitHub repository: https://github.com/samansalike/DR-thermo

## 1 Introduction

Thermodynamic analysis of biochemical reaction systems provides a means to analyze the equilibrium states of the reactions. Further, a thermodynamic analysis is useful to assign directionalities, evaluate feasible pathways, and to integrate metabolomics data into mathematical models (Ataman and Hatzimanikatis, 2015). Applications of biothermodynamics to evaluate pathway feasibility have been explored in several areas of systems biology (Donnelly and Wolfe, 1986;Eaton and Chapman, 1992;Dolfing, 2000). For instance, thermodynamic properties have been coupled with the metabolic flux analysis (MFA) to limit fluxes through futile cycles while still allowing fluxes through feasible pathways (Henry *et al*., 2007). Incorporation of thermodynamic properties has broadened the applicability of MFA methods without adversely affecting their accuracy (Garg *et al*., 2010). The use of such tools has helped elucidate genotype-phenotype relationships. Further, it has aided metabolic and genetic engineering efforts to produce commercially desirable compounds via heterologous routes (Long *et al*., 2015).

Among the thermodynamic properties, the Gibbs free energy of a reaction is of primary importance because it permits the quantification of the degree of thermodynamic favorability of the given reaction (Goldberg, 2014). Since biochemical reactions occur in aqueous solutions and involve the release or uptake of protons, the standard ‘transformed’ Gibbs energy of a reaction depends on the ionic interactions and can be calculated from the apparent equilibrium constant, *K’*. Currently, empirical data on apparent equilibrium constants can be obtained from databases such as the National Institute of Standards and Technology (NIST)’s Thermodynamics of Enzyme-Catalyzed Reactions Database (TECRDB) (also labelled as NIST-TECRDB) (Goldberg *et al*., 2004, 2007) which contains a comprehensive list of *K’* values for over four hundred reactions under different conditions. Moreover, since Gibbs energies of a reaction can also be computed from individual formation energies, several studies have also been dedicated to determine formation energies and have generated commendable data banks (Alberty, 1998, 2005; Thauer *et al*., 1977; Thauer, 1998; Wagman *et al*., 1982). However, thermodynamic analysis based entirely on such experimental databases is limited to small-scale systems or a subsections of genome-scale systems (Jankowski *et al*., 2008). Consequently, several estimation methods have been formulated over the years to bridge this gap, the most prevalent of which is the group contribution method developed by Mavrovouniotis (Mavrovouniotis, 1990, 1991). This method in due course was greatly improved upon by several researchers (Jankowski *et al*., 2008;Noor *et al*., 2012, 2013; Du *et al*., 2018). The group contribution method is based on the idea that the overall formation energy of a compound can be approximated as a linear function of the ‘contributions’ of its functional groups. The values of these group contributions are estimated using by a linear regression on the data set consisting of experimentally determined standard Gibbs free energies of reaction and formation. Such empirical prediction methods have successfully been used to study aromatic amino acid pathways (Hatzimanikatis *et al*., 2005), glycolysis (Maskow and von Stockar, 2005), genome-scale model of E. Coli (Feist *et al*., 2007; Henry *et al*., 2006, 2007) biodegradation pathways (Finley *et al*., 2009), and the metabolic network of G. sulfurreducens (Garg *et al*., 2010).

However, inaccuracies in experimental data can lead to inconsistencies when different databases are combined to calculate property values for a given reaction (Goldberg, 2014). Such inconsistencies are also reflected in the estimated values for missing Gibbs energies of reaction and formation which are computed from the available measurements. Often, these estimates, as well as the measurements of Gibbs energies, violate the first law of thermodynamics because the overall Gibbs energy of a reaction that takes place in more than one step does not correspond with the sum of all standard Gibbs energies of intermediate steps under the same conditions. As a result, futile cycles can have a nonzero change in Gibbs energy (Noor *et al*., 2013). Moreover, the effects of ionic interactions on the experimental data have to be normalized in order to obtain linear relationships between all variables (Alberty, 2005). This involves using the reported measurements of pH, ionic strengths and proton dissociation constants, which are often erroneous or missing altogether. For example, Maskow and von Stockar, 2005 found that feasible pathways can be falsely labelled as infeasible and vice versa without careful consideration of ionic strength of the solution, uncertainty in thermodynamic data and cell pH in studying the thermodynamic feasibility of the lactic acid fermentation pathway.

Such difficulties arising due to measurement errors are not unique to systems biology. For instance, plant data obtained in chemical and bio-tech industries are often corrupted by measurement errors which undermine the quality of such data-driven methods. In such scenarios, data reconciliation (DR) as a technique has been used for an accurate and consistent estimation of process parameters and variables from measurements (Dabros *et al*., 2009; Narasimhan and Jordache, 1999; Hodouin, 2010). The DR based methods exploit the redundancies arising due to physical constraints such as the closure of mass and energy balance equations to improve the estimates of the underlying variables (Narasimhan and Jordache, 1999). The aim of the DR methods is to obtain accurate estimates of the variables from experimental data such that they are consistent with the physical constraints imposed (Narasimhan and Jordache, 1999). Since physical laws of mass conservation and thermodynamics also govern biosynthetic transformations in biological systems (Lewis *et al*., 2012), we show that a similar approach can be employed to obtain consistent estimates of Gibbs free energies of reaction and formation.

In this work, we propose a framework that employs the data reconciliation approach to adjust available measurements such that thermodynamic laws are satisfied. It will be shown that the proposed DR approach can be used to reconcile all measured Gibbs energies and estimate the missing Gibbs energies under certain conditions when a subset of measurements is available. Further, the DR framework is extended to estimate group contributions. Moreover, we also provide guidelines for the curation of the reconciled data and the design of new experiments for an improved estimation of missing Gibbs free energies. To illustrate the applicability of the framework, we use a subset of reactions from the NIST-TECR database that only contain compounds whose formation energy value is available from the Alberty (2005) table.

## 2 Approach

The DR framework is shown in Figure 1 for estimating Gibbs free energies. In the reconciliation step, the DR framework requires three inputs: (i) measured variables (the standard ‘transformed’ Gibbs free energies of reaction and formation), (ii) a priori information such as stoichiometric matrix, group definitions, and (iii) constraints between all the variables. The experimental standard ‘transformed’ Gibbs energies are converted to standard Gibbs energies by applying the inverse Legendre transform, using the proton dissociation constants and the values of the controlled variables such as temperature, *pH* and ionic strength. The reconciled Gibbs free energies are then estimated by minimizing the sum squared difference between each measurement and the corresponding estimate, subject to thermodynamic constraints. The missing Gibbs energies of reaction and formation are estimated by using the reconciled estimates of the measurements and the constraint equations representing their relationships. Since the thermodynamic constraints are linear, this leads to a linear DR problem which can be solved analytically. The DR framework provides thermodynamically consistent estimates of Gibbs free energies. Then, these reconciled values of Gibbs free energies of reaction, formation and groups can be curated in a database.

In the prediction step, the curated data can be used to estimate any new unknown Gibbs free energies of reaction. If the Gibbs free energies of formation for all the reactants taking part in a given reaction are available in the curated database, the thermodynamic constraints can be directly used to predict the Gibbs free energy of the given reaction, often labelled as the standard method. On the other hand, if the Gibbs energies of formations for only a subset of reactants are available in the curated database, the component contribution method can be used to predict the reaction Gibbs free energies using the available Gibbs energies of formation and the group contributions (Noor *et al*., 2013). Further, the information of temperature can also be incorporated (Du *et al*., 2018). This work deals with the first step of data reconciliation formulation to obtain the thermodynamically consistent estimates of Gibbs free energies.

## 3 Methods

In this section, we delineate the data reconciliation approach to consistently estimate the Gibbs free energies of reaction, formation, and group contributions from experimental data.

**Fig.1.**
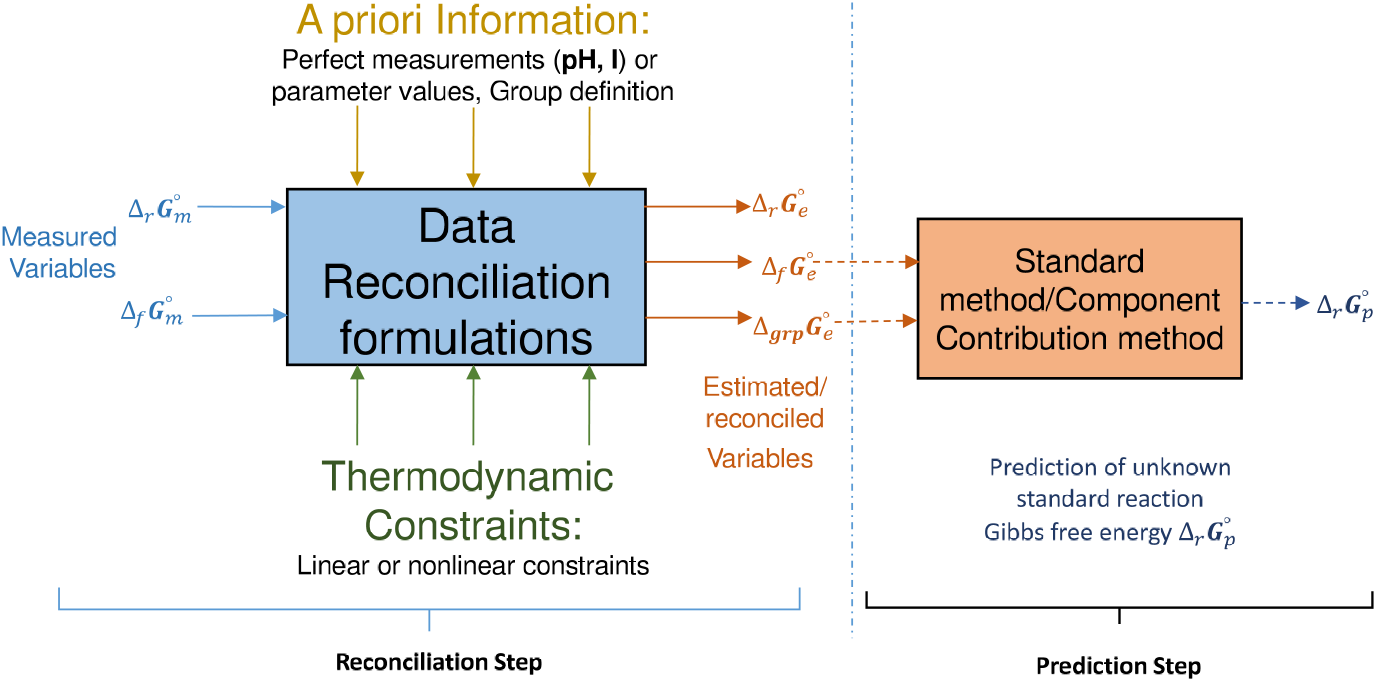
Data Reconciliation Framework consisting of two parts: (i) Reconciliation Step: Thermodynamically consistent estimates of Gibbs free energies of reaction, component formation and groups are obtained and the reconciled estimates are curated for the prediction step, and (ii) Prediction step: The Gibbs free energies of unknown reactions are predicted using the reconciled formation and/or Groups Gibbs free energies and standard method or component contribution method from the curated database.

### 3.1 Data reconciliation: Imposing thermodynamic constraints for estimating standard Gibbs free energies

Consider a biochemical reaction network with *n* reactions between *m* compounds. If the vectors 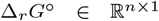 and 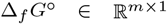 represent the standard Gibbs energies of reaction and formation, respectively^1^. The stoichiometric matrix 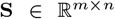 describes the metabolic reactions taking place in the system. Each column in S represents the stoichiometric coefficients of a reaction with negative and positive values representing the reactants and products respectively. At the standard conditions, the thermodynamic law leads to the following relationship (Alberty, 2005):

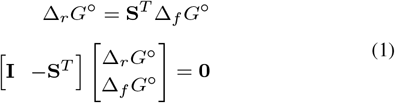

Eq. (1) is violated due to the presence of measurement errors in the measured Gibbs energies, Δ*_r_G°_m_* and Δ*_f_G°_m_*. Hence, the objective is to estimate these energies such that they do not violate the thermodynamic relationships in Eq. (1). Note that there are *n* relationships in Eq. (1) for the *n* reactions. Often, it is assumed that the measurement errors follow a multivariate normal distribution with zero mean and a co-variance matrix Σ. Then, the DR problem can be formulated as a constrained weighted least–squares optimization problem as follows:

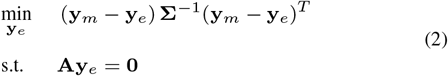

with 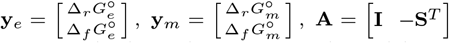, where the subscripts m and e denote the experimental data and the corresponding reconciled estimates, respectively. In the objective function, total (*n+m*) Gibbs free energies are the reconciled (or estimated) such that they satisfy the *n* thermodynamic constraints in Eq. (1). Each diagonal element of Σ represents the variance of the measurement error in a variable. Thus, higher weights are given to more accurate measurements. In practice, these variance values can be estimated by repeated measurements or the experimentalist can assign values based on their knowledge. Otherwise, a weight of one can be assigned. It should be noted that the standard transformed Gibbs energies Δ*_r_G’°* (computed fromapparent equilibrium constants) include the potential of *H*^+^ as a natural variable. This is the effect normalized by the inverse Legendre transform while converting the standard ‘transformed’ Gibbs energies to standard ‘chemical’ Gibbs energies that are used to impose thermodynamic constraints on estimates using Eq. 1 (see Section 1 of the supplementary material for details on the inverse Legendre transform).

The constrained optimization problem in Eq. (2) is a quadratic optimization problem. Hence, an analytical solution of the optimization problem Eq. (2) for **y***_e_* can be computed using the first order optimality conditions as (Narasimhan and Jordache, 1999):

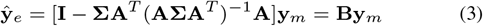

Then, the estimated 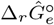 and 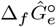 can be obtained in terms of the measured values 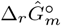 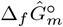 as follows:

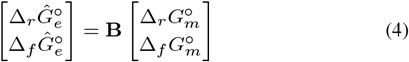

Further, note that the estimates in Eq. (4) are normally distributed with the expected value and co-variance matrix given as follows (Narasimhan and Jordache, 1999):

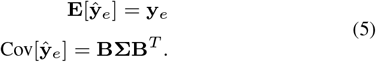

The characteristics of the estimates in Eq. (5) are useful in providing bounds in thermodynamics-based metabolic flux analysis or metabolic network thermodynamics (Henry *et al*., 2007).

#### 3.1.1 Partially measured reaction systems

Often, the Gibbs free energies of all reactions and compound formation in a biochemical reaction network are not measured. Hence, a subset of Gibbs free energies measurements is only available. In such cases, the linear DR framework can be extended to reconcile the measured Gibbs free energies and to estimate the unmeasured ones under certain conditions. In this situation, this DR problem is decomposed into two sub-problems. In the first sub-problem, the approach mentioned based on the fully measured Gibbs free energies (Section 3.1) has to be modified such that a reduced set of constraints between the measured Gibbs free energies is obtained. Then, the reconciled estimates of the measured Gibbs free energies are obtained subject to the reduced set of constraints. In the second step, the thermodynamic constraints are used to estimate the unmeasured Gibbs free energies from the reconciled ones obtained in the first step. To obtain a reduced set of constraints in the first sub-problem, the constraints in Eq. (1) can be decomposed into the measured (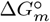) and unmeasured Gibbs free energies (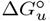) as follows:

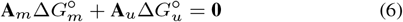

where the matrix **A_m_** consists of the columns of [**I**–**S**^*T*^] corresponding to 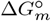 while the matrix **A_u_** consists of the columns of [**I** – **S**^*T*^] corresponding to 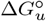. In practice, the Gibbs free energies of reactions and a subset of the Gibbs free energies of formation are measured. For a reaction system involving *n* reactions and *m* species, if only *p* Gibbs free energies of formation (< *m*) are not measured. Then, Eq. (6) can be written as follows

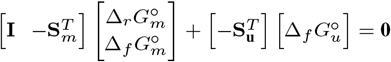

where 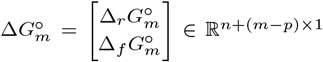 and 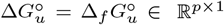 are vectors representing the measured reaction and formation energies and the unmeasured formation energies. The formulation in Eq. (6) is also applicable when the Gibbs energies of reactions are not measured. Similarly, **S**_*m*_ and **S**_*u*_ are the stoichiometric matrices corresponding to the measured and unmeasured formation energies. The contribution of the unmeasured Gibbs free energies can be eliminated by constructing a projection matrix **P** of dimension (*q* × *n*) such that **PA_u_** = **0**. Then, by pre-multiplying the constraints in Eq. (6), the reduced set of constraints corresponding to the measured Gibbs free energies are obtained as follows:

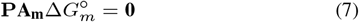

Note that there are only *q* < *n* constraints in the reduced set. The details for the computation of project matrix are given in Section 4 of the supplementary material. The resulting DR problem can be formulated as follows:

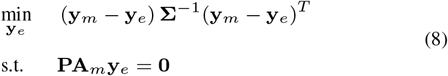

with 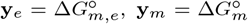. The reconciled estimates, 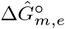, can be computed using the solution in Eq. (3) with **A** = **PA**_*m*_.

In the second step, the unmeasured Gibbs free energies can be estimated using a least-squares solution as follows:

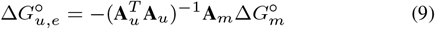

Asufficient number ofGibbs free energies have to be measured to apply the approach mentioned based on the partial measurements. In construction of **P** for this approach, it is assumed that the *q* rows of **P** are independent and *q* = *n* – *p*. Hence, all the *p* unmeasured formation energies can be estimated. However, even if the *q* rows are not independent, only a subset of the unmeasured formation energies can be estimated. The analysis related to solvability and estimability of the Gibbs free energies for a given set of Gibbs free energies measurements is discussed in Section 3.3.

### 3.2 Estimating group contributions

Improving the accuracy of the group contribution methods is of prime importance due to their high coverage and applicability to the majority of biochemical reaction networks (Noor *et al*. (2013)). However, in addition to the effect of measurement errors, inaccuracies in these estimates arise due to the simplifying assumption that the group contributions are additive. This is referred as modelling error due to result of the approximation 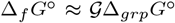, where 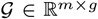 is the group incidence matrix in which each row represents the decomposition of each compound in **S** into to a predefined set of *g* structural subgroups and the vector 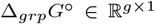 consists the Gibbs free energies of different groups as its elements. Since the relationships between the Gibbs energies of groups and species’ formation are empirical in nature, these relationships cannot be included as constraints. However, an improved optimization problem can be formulated such that measurement and modelling errors are simultaneously minimized without violating the thermodynamic constraints. This is achieved by using the reconciled estimates of formation energies to estimate group contributions by modifying the optimization problem. This optimization problem extends the formulation in Eq. (2) to include the information of the group contribution method, with 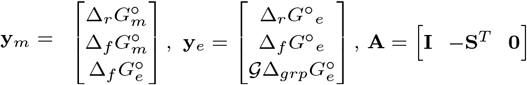. Here, Δ_*f*_*G*°_*e*_ is included in both **y**_*e*_ aswell as **y**_*m*_. The estimation of group contributions for a rank-deficient matrix 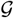 is given in Section 7 of the supporting material.

### 3.3 Observability and redundancy analysis of measurements

For a given set of measured Gibbs free energies of reaction and formation, it is important to answer the following two questions: (Q1) Is it possible to obtain thermodynamically consist reconciled estimates for the given set of measurements? and (Q2) Is it possible to estimate unmeasured missing Gibbs free energies from the measured ones in partially measured systems? The DR framework can provide answers for these questions using the concepts of redundancy (for Q1) and observability (for Q2). The observabiity and redundancy analysis is performed as a part of the DR approach. For a reaction system with *n* reactions and *m* compounds, if an unmeasured Gibbs free energy can be estimated uniquely from the measured Gibbs free energies and the constraints, then the unmeasured Gibbs free energy is observable. On the other hand, if measurement of a Gibbs free energy is removed from the measured ones, and it can be still uniquely estimated from the rest of the measured energies, then the measured one is redundant and this property is referred as redundancy (See Section 5 of the supporting materials for details). For a partially measured reaction systems, if the *s* rows of **P** are independent with *s* < *q*, then the observability analysis allows us to find out which unmeasured Gibbs free energies can be estimated from the measured ones. Let us divide the unmeasured Gibbs free energies 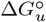 into two parts as follows: 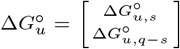. Then, the following relationships can be derived
between 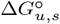 corresponding to the *s* independent rows of the projection matrix **P**, 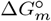 and the remaining *q* – *s* unmeasured Gibbs free energies 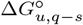 as follows:

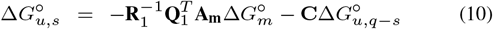

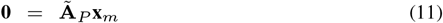

where **R**_1_, **Q**_1_, 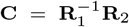, and 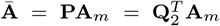 are the matrices obtained via the QR decomposition of **A**_*u*_. The details of derivation and the matrices are given in Section 5 of the supporting materials. In Eq. (11), the matrices **C** and 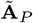 contain all the necessary information to classify Gibbs free energies into the observable and redundant variables. If a row of **C** has zero elements only, then the corresponding unmeasured Gibbs free energies can be estimated uniquely from 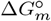 and it is observable. On the other hand, if a row of **C** contains at least one non-zero element, then the corresponding Gibbs free energy is unobservable because its estimates depends on the values of the (*q* – *s*) unmeasured Gibbs free energies as it can be seen in Eq. (11). Further, the (*q* – *s*) unmeasured energies are also unobservable because their values have to be specified. If a column of 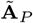 has zero elements only, then the corresponding measured Gibbs free energy is not the redundant variable. In other words, that energy can not be reconciled as it cannot be indirectly estimated from the other measured energies and the reduced set of constraints (Narasimhan and Jordache (1999)).

## 4 Results and Discussion

**Published data**: We consider a test set of 87 different metabolic reactions from the NIST-TECR database to illustrate the data reconciliation framework. The measured formation energies of the 84 compounds involved were obtained from (Alberty, 2005). The relevant data are provided in Section 9 of the supplementary material. The measurements of the apparent equilibrium constants for many of these reactions are available at different conditions. The inverse Legendre transformation was applied to calculate the standard Gibbs free energies of these reactions.

**Simulated data**: To assess the accuracy of the DR approach, the true values of the available thermodynamic data have to be known. Thus, the standard reaction and formation energy estimates obtained after reconciling the published values were assumed to be “true” parameter values for the simulated data. Then, noisy Gibbs free energy measurements were generated by adding the Gaussian errors with zero mean and variance of *α* percentage of the “true” value of each measurement where *α* is a positive constant. These simulated data were used to reconcile the Gibbs free energies using the DR approach and the regression methods.

### 4.1 Comparison of data reconciliation with the linear regression

The DR method and regression are applied to the published experimental Gibbs free energy data. The results are shown in Figure 2. It can be observed from Figure 2 a) that the published experimental Gibbs free energy data are not thermodynamically consistent, i.e. they do not satisfy constraints in Eq. (1). Thermodynamic consistency here is the deviation of the 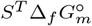 value from the corresponding measured 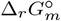 value (vertical distance from the 45^°^ line). Since the erroneous experimental data were combined from different sources, the average constraint error in terms of the root-means-squares-error (RMSE) is 22.63 kJ mol^−1^. Under the assumption that the uncertainty is only due to measurement errors in apparent equilibrium constant data, and not in the normalization of ionic effects, the DR method as applied to the published data. The resulting reconciled Gibbs free energies after adjusting both 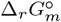 and 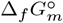 were consistent, with zero constraint errors as shown in Figure 2 a).

**Fig. 2.**
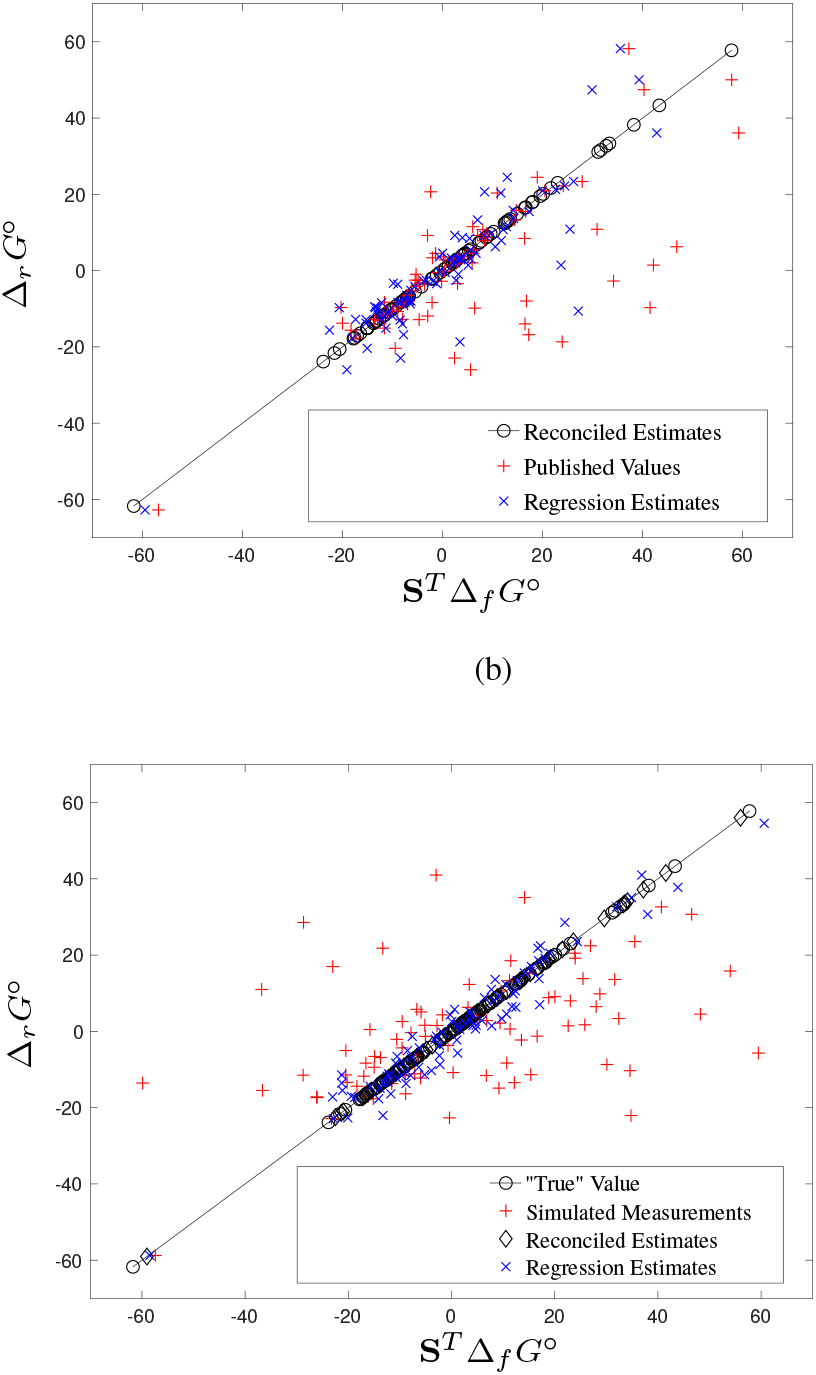
(a) Comparison of the extent of violations of the thermodynamic constraints (Eq. (1)) in the published, regression estimates and the reconciled estimates for standard Gibbs free energies. The average constraint error per reaction in terms of the RMSE values are 22.63 kJmol^-1^ and 7.34 kJmol^-1^ are for the published data and the regression estimates, respectively. This error is virtually zero in the reconciled estimates. (b) Comparison of the estimates for the simulated data with 10% noise with RMSE of 24.16 kJmol^-1^, similar to published data. The resulting reconciled estimates are consistent and have an average deviation of 2.94 kJ mol ^-1^ from the “true” values in comparison to 4.64 kJ mol^-1^ in the regression estimates.

The linear regression between the experimental data 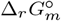 and 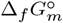 with the model equations being the constraints (Eq. 1) is fitted (see Section 2 for more details). In the linear regression, it is assumed that the formation energy measurements do not contain any measurement errors. However, due to the measurement errors in the experimental data in the training data and only 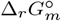 error is minimized without imposing the thermodynamic constraints. This approach leads to thermodynamic inconsistencies in the estimates which cannot be avoided. Hence, the average constraint error in terms of the RMSE is 7.34 kJ mol^-1^ in the linear regression estimates of the measured data. The accuracy of an estimation method is determined by the deviations of the estimates from the true values. In this study, this error is referred to as a prediction error (The details on the constraint error and prediction error are provided in Section 8 of the supplementary material). To demonstrate the utility of the DR approach, we have applied the DR and linear regression method to the simulated data. As shown in Table 1, even for the *α* value as high as 30 %, the average deviations from the “true” values in the case of linear DR estimates of the simulated measurements are nearly half of those of the linear regression estimates. For the simulated data with *α* = 10%, the estimates and confidence intervals are given in Section 10 of the supporting information.

**Table 1.** Comparison of linear data reconciliation and linear regression for simulated data. The errors computed as an RMSE in kJ mol^-1^

| $\alpha$ | Average<br>constraint error<br>(Reconciliation) | Average<br>constraint error<br>(Regression) | Average<br>prediction error<br>(Reconciliation) | Average<br>prediction error<br>(Regression) |
| --- | --- | --- | --- | --- |
| 1% | $2.86 \times 10^{-12}$ | 1.4613 | 0.949 | 1.3327 |
| 5% | $2.74 \times 10^{-14}$ | 4.2721 | 1.9028 | 3.8284 |
| 10% | $1.19 \times 10^{-12}$ | 5.8771 | 2.947 | 4.639 |
| 20% | $6.59 \times 10^{-12}$ | 9.0241 | 3.4290 | 7.0352 |
| 30% | $1.24 \times 10^{-11}$ | 9.9850 | 4.3306 | 8.1603 |

The main use of the DR method is in estimating missing Gibbs energies and reconciling estimates from measured Gibbs free energies in partially measured systems. An observability analysis for the entire set of reactions on the NIST-TECR database of nearly 4000 reactions between 566 compounds. It was found that the formation energy estimates of 77 compounds can be obtained entirely from the available experimental data on the formation energies (of 117 compounds obtained from Alberty (2005)) and the reaction equilibrium constants for 134 reactions. (See the reconciled values for 77 compounds in Section 6 of the supporting information.) To evaluate the regression and DR approach, we used subsampling based cross-validation using the simulated data (with *α* = 10). To ensure that only the observable Gibbs energies are sampled out, the observability analysis was performed for the matrix [**I** – **S**^*T*^] and the 87 observable Gibbs energies of reaction and formation were identified. For generating a sample, thirty observable Gibbs energies were randomly sampled out as a validation set. The remaining data was used to train the regression model and the estimates for the validation set were then determined. The method for linear DR for partially measured measured reaction systems in Section 3.1.1 is applied to obtain the reconciled and estimates of the unmeasured Gibbs free energies in a validation set. The process of sub-sampling and estimation was repeated several times with increasing number of sub-samples from 10 to 500. After each step of sub-sampling and estimation, the average cross-validation error was then computed as RMSE of the sub-samples for the estimates obtained by the linear regression and DR method. The results are shown in Figure 3 as a plot between the resulting RMSE of sub-samples and the number of subsamples. The average cross validation error for regression was higher than that of the DR approach at the low number of sub-samples. In fact, this error was initially very high, but approached that of reconciliation as the number of iterations increased beyond 500. The validation error for reconciliation remained roughly constant irrespective of the sub-sample being validated. This suggests that regression is prone to over-fitting and the accuracy of the method is sensitive to the quality of training data and the exact set of Gibbs energies that are required to be estimated. The DR avoids this by using reconciled estimates of measurements with zero constraint error to estimate the missing Gibbs free energies.

**Fig. 3.**
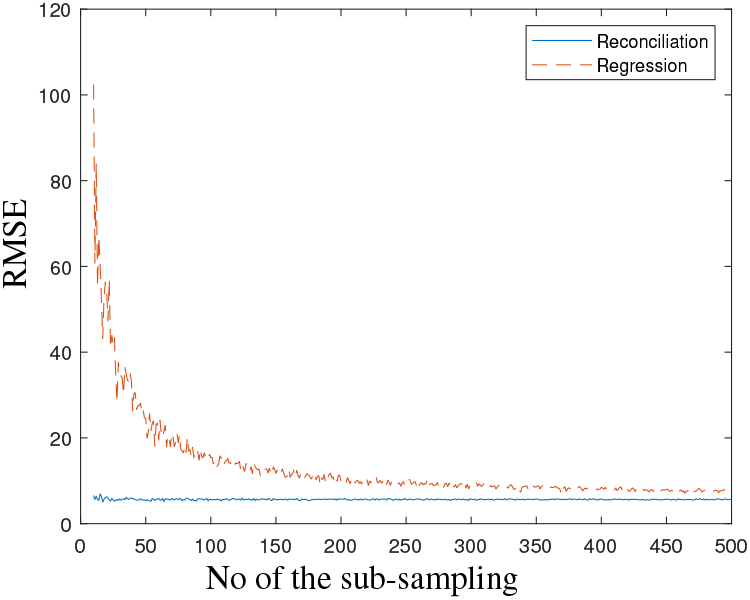
The results of sub-sampling cross validation of 30 Gibbs energies. The validation errors (RMSE) for the regression was higher than those of the DR approach.

### 4.2 Improved estimation of group contributions

The compounds in the data set considered were decomposed into thirty groups and the group incidence matrix *G* was constructed. The group decomposition was carried out based on the work of Noor *et al*., 2013 and the same group definitions were used (see Section 10 of the supporting material for details). For simulated measurements with *α* = 10, the group contributions were estimated using the DR approach in Section 3.2. The constraint error of the DR estimates quantified here by the average squared difference between Δ_*r*_*G*° and **S**^*T*^*G* · Δ_*grp*_*G*° was lower (RMSE of 3.77 kJ/mol) in comparison to the one obtained by the regression approach (RMSE of 5.83 kJ/mol) as shown in Figure 4. The residual inconsistency present in the reconciled estimates can be attributed to the modelling error due to the group additivity assumption. The group contribution estimates of Δ_*r*_*G*° also had slightly lesser prediction errors when the reconciliation approach was used (RMSE of 5.35 kJ/mol compared to 6.05 kJ/mol using regression). However, the thermodynamic constraints in (1) are satisfied.

**Fig. 4.**
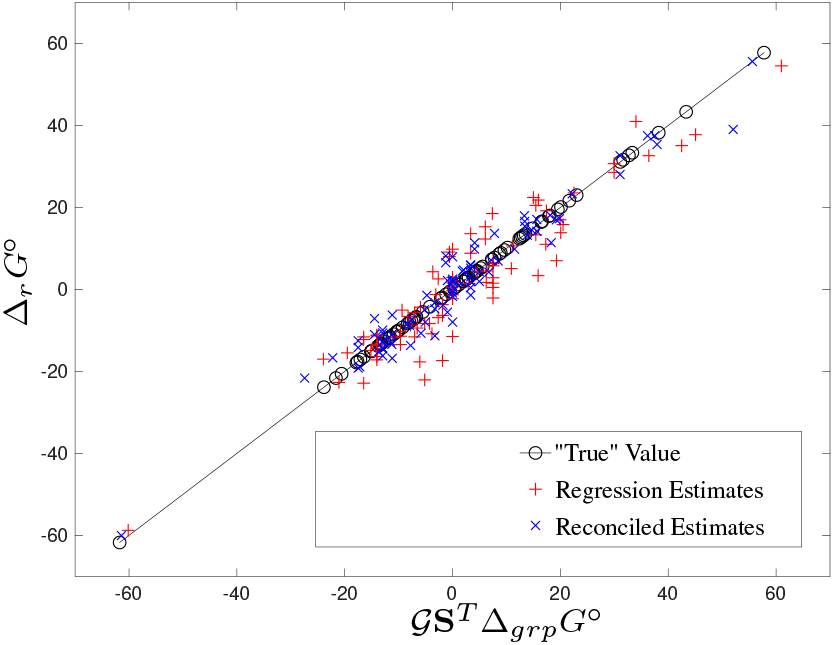
Results of the modified group contribution showing that the reconciled estimates are more thermodynamically consistent (constraint error of 3.77 kJ/mol) than the regression estimates (constraint error of 5.83 kJ/mol). Morever, the reconciled estimates of group contributions yield reaction energies that are closer to the “true” values. (RMSE of 5.35 kJ/mol compared to 6.05 kJ/mol for the Pseudoisomer group contribution (PGC)

## 5 Curation of data and Design of experiments

For an existing database, if new Gibbs free energy measurements are available for a given reaction or species from the database, its redundancy is increased. The DR framework can thus accommodate systematic revisions of a large ‘global’ database such as TECRDB based on the addition of new experimental data replicates, contributing to increased accuracy of the reconciled estimates. Moreover, in curating a ‘global’ database, the unobservable Gibbs energies of reaction and formation can be identified and used as salient targets for designing new experiments. For instance, for the 473 unique reactions on the NIST-TECR databse, experimental thermodynamic data for formation energies is only available for 117 of the 566 compounds involved in the reactions (from the (Alberty, 2005) table). Due to lack of observability, not all of the remaining formation energies can be estimated. In fact, using the data-reconciliation framework, it was found that formation energies of 77 compounds are observable and can be estimated using the DR approach for partially measured systems without requiring any additional information (see Section 6 for the details of estimates). However, for 150 of the unmeasured compounds, their formation energies are required to be specified to estimate the rest. Thus, conducting experiments to find the formation energies of these compounds can effectively ‘complete’ the database. When experimental data is not available for unobservable Gibbs energies, the group contribution method using the available measurements within the database can be can be used to impute estimates for these variables, albeit at reduced accuracy. Other non-empirical methods (e.g. Jinich *et al*. (2014)) can also be used for this imputation.

While curating databases and adding new information, an important distinction should be made between a ‘global’ database and a new experimental dataset. When a new set of reactions is considered, the data should be collated with the ‘global’ database when the size of the two sets are comparable. This is because when the DR is performed on a collated set of reactions, new reconciled estimates are obtained for both ‘global’ and the new experimental data. If the new experimental data are not large enough, modifying the values in the ‘global’ database based on limited new experimental data can lead to inaccuracies. The reconciled estimates from the global set can only be used to predict missing information in the the new set using the standard method or the component contribution method. If the prediction set is partially measured, the available measurements can only be reconciled using the constraints within the set. However, When the size of the new set is large enough, the two sets can be collated to improve estimation of new Gibbs energies and the coverage of the database.

## 6 Conclusion

In this study, the data reconciliation (DR) framework has been proposed for obtaining thermodynamically consistent estimates of Gibbs free energies from experimental data. The DR framework has been extended to estimate group contributions. Further, for a partially measured system, using both published and simulated Gibbs energies data, it has been shown that the proposed methods can be used to reconcile available Gibbs free energy measurements and provide consistent estimates for the missing ones that have better accuracy than linear regression.

Given a set of reactions and all the available experimental thermodynamic data on Gibbs energies, the DR framework can thus be used to reduce experimental errors in the data by reconciling all measurements, classify missing Gibbs energies as observable or unobservable, and calculate consistent estimates for the observerable Gibbs free energies. This a priori analysis can also aid in designing new experiments in a systematic manner and help obtain thermondynamically consistent estimates of all Gibbs free energies. The DR methods can also improve the curation of data banks from different sources to increase the scope of estimation and applicability of the available information, potentially becoming an invaluable tool for the thermodynamic analysis of biochemical systems.

## Supporting information

## Acknowledgement

We would like to thank Noor *et al*. (2013) for the open sourced component contribution framework on GitHub (https://github.com/eladnoor/component-contribution).

## Footnotes

1 The standard condition in biochemical systems corresponds to Pressure, *P* = 1 bar, *pH* = 7, and *pMg* = 3, and the ionic strength, *I* = 0 or *I* = 0.25 M. Often, *I* = 0.25 is defined as near physiological conditions (Alberty, 1994)

