## Supplementary material for "Thermodynamically Consistent Estimation of Gibbs Free Energy from Data: Data Reconciliation Approach"

[3] Initiative for Biological Systems Engineering (IBSE)

[4] Robert Bosch Centre for Data Science and Artificial Intelligence (RBC-DSAI),

India Indian Institute of Technology Madras, Chennai –600036, India

**3 Theoretical basis of data reconciliation** **4**

**4 Construction of a projection matrix using QR decomposition** **4**

**5 Observability and Redundancy** **5**

**6 Analysis of NIST-TECR Database for Observability and Redundancy** **6**

**7 Estimation of group contributions when $\mathcal{G}$ is rank deficient** **9**

**8 Calculation of errors** **9**

### 1 Inverse Legendre transform

The inverse Legendre transform is used to convert apparent (or transformed) Gibbs energy of formation $\Delta_f G'^{\circ}$ to its standard Gibbs energy of formation $\Delta_f G^{\circ}$. The main idea here is that the thermodynamics of biochemical systems is better discussed in terms of compounds, or an ensemble of species such as ATP as opposed to species like $HATP^{-3}$, $ATP^{-4}$, in order to describe their metabolism [2]. Thus, when measurements of *K'* are made at equilibrium at a constant pH, the standard 'transformed' Gibbs energy, $\Delta_f G'^{\circ}$, corresponding to this measurement at equilibrium is defined at a specified temperature $T$, pressure $P$, and $pH$ and has the chemical potential of $H^+$ as a natural variable. For a species *j*, the transformed Gibbs free energy at the $pH$ and ionic strength $I$ can be given as [1]:

$$\Delta_f G_j'^{\circ}(pH, I) = \Delta_f G_j^{\circ} + (N_H RT \ln 10)pH - \left(\frac{2.91482({z_j}^2 - N_H)I^{1/2}}{1 + 1.6I^{1/2}}\right) \tag{1}$$

where $I$ is the ionic strength of the solution which is dependent on temperature, the charge of the species $j$ (denoted as $z_j$), and the number of hydrogen atoms in the species $j$(denoted as $N_H$). For a reactant *i*, which is an ensemble of species $j$, the standard 'transformed Gibbs energy can be given by:

$$\Delta_f G_i'^{\circ} = -RT \ln \sum_j e^{-\Delta_f G_j'^{\circ}/RT} \tag{2}$$

The analysis of biochemical reaction systems using Eqs. (1) and (2) can be reduced by aggregating species with the same chemical potentials, and making thermodynamic calculations with isomer groups. However, when pH is specified, groups have the same values of the transformed chemical potential, and the species in a group are referred to as pseudoisomers. Isomers have the same chemical potential at equilibrium while pseudoisomers have the same transformed chemical potential at equilibrium at a specified pH [2]. The pseudoisomers represent different protonated species of reactants and the difference between two consecutive pseudoisomers is a function of the corresponding $pK_a$ value. Thus, we can express any $\Delta_f G_i^{\circ}(n)$ as a function of a single reference pseudoisomer $\Delta_f G_i^{\circ}(m)$ and the list of $pK_a$ values using [5]:

$$\Delta_f G^{\circ}(n) = \Delta_f G^{\circ}(m) - RT \ln 10 \sum_{i=m+1}^{n} pK_a(i) \tag{3}$$

Then, by combining Eqs. (1), (2) and (3), the Legendre transformed which is used to define the standard 'transformed' Gibbs energy of a reactant is given by::

$$\Delta_f G'^{\circ}(n) = \Delta_f G^{\circ}(m) + RT \ln 10 \left(m \cdot pH + \sum_{i=m+1}^{n} pH - pK_a(i)\right) - \frac{2.91482({z_n}^2 - n)I^{1/2}}{1 + 1.6I^{1/2}} \tag{4}$$

where $z_n$ is the net charge of species with *n* hydrogens. Eq. (4) can be re-arranged to calculate the standard Gibbs energies from the standard 'transformed' Gibbs energies. thus performing the function of an 'inverse' Legendre transform.

### 2 Regression Model

The empirical prediction methods currently used are based on the calculation of compound formation energies by a linear regression on the Gibbs energy of reactions obtained from measurements of equilibrium constants. As discussed in Section 1, the apparent Gibbs energies of reactions are first converted to standard Gibbs free energies using the inverse Legendre transform. This method has been revised and improved over the years, with the highest accuracy achieved in the work of [5] by accounting for $\boldsymbol{pH}$ and $\boldsymbol{I}$. Further, It has been shown that it is possible to combine more accurate reactant contribution method (PRC) with the less accurate group contribution method (PGC) to achieve better results and wide coverage [6]. The PRC and PGC methods are compared with the data reconciliation approach in this study and are described briefly in this section. Note that the prediction step of data reconciliation framework will use the standard method or PRC-PGC based method to predict the Gibbs free energy of an unknown reaction using the reconciled estimates.

#### 2.1 Pseudoisomer reactant contribution (PRC) method

Consider a system of $\boldsymbol{n}$ reactions between $\boldsymbol{m}$ compounds with the stoichiometric matrix given by $\boldsymbol{S} \in \mathbb{R}^{\boldsymbol{m}\times\boldsymbol{n}}$. The formation energy estimates, represented by the vector $\boldsymbol{\Delta_f G^\circ_{est}} \in \mathbb{R}^{\boldsymbol{m}\times 1}$, can be obtained from the available measurements of reaction Gibbs energies, given by $\boldsymbol{\Delta_r G^\circ_{obs}} \in \mathbb{R}^{\boldsymbol{n}\times 1}$ using a regression model as follows:

$$\boldsymbol{\Delta_f G^\circ_{est}} = (\mathbf{S^T})^+ \cdot \boldsymbol{\Delta_r G^\circ_{obs}}$$

Here, $(\mathbf{S^T})^+$ is the Moore–Penrose pseudo-inverse of the stoichiometric matrix. The estimates obtained for the Gibbs free energies of formation are then used to predict the reaction energies of a new reaction if its stoichiometric vector is in the range space of $\boldsymbol{S^T}$. Due to this limitation, the coverage of this method is low.

#### 2.2 Pseudoisomer group contribution (PGC) method

In this approach, the total formation energy of a reactant is modelled as a sum of the contributions of a set of structural groups present in it. Thus, each compound in $\boldsymbol{S}$ is decomposed to a predefined set of $\boldsymbol{g}$ structural subgroups and each decomposition is represented by a row of the group incidence matrix $\boldsymbol{\mathcal{G}} \in \mathbb{R}^{\boldsymbol{m}\times\boldsymbol{g}}$. Then, an approximate regression between the Gibbs free energies of a reaction and groups can be written as:

$$\boldsymbol{\Delta_r G^\circ} \approx \mathbf{S^T}\boldsymbol{\mathcal{G}} \cdot \boldsymbol{\Delta_{grp} G^\circ}$$

where $\boldsymbol{\Delta_{grp} G^\circ} \in \mathbb{R}^{\boldsymbol{g}\times 1}$ is a vector representing the Gibbs free energies of group contributions. Then, the Gibbs free energies of groups can be obtained from the experimental data as follows:

$$\boldsymbol{\Delta_{grp} G^\circ_{est}} = (\mathbf{S^T}\boldsymbol{\mathcal{G}})^+ \cdot \boldsymbol{\Delta_r G^\circ_{obs}}$$

Then, during the prediction stage While this method gives increased coverage, its accuracy is low. This is due to the assumption that the overall formation energy is a linear sum of the individual group contributions.

#### 2.3 Effect of measurement error on regression estimates

The values of $\boldsymbol{\Delta_r G^\circ_{obs}}$ are obtained experimentally and are thus subject to measurement noise. Due to this, the regression model takes the form:

$$\boldsymbol{\Delta_r G^\circ_{obs}} = \mathbf{S^T} \cdot \boldsymbol{\Delta_f G^\circ} + \boldsymbol{\varepsilon}$$

Here, $\boldsymbol{\varepsilon}$ represents the random error. Since the regression model is fitted on empirical data, this error is manifested in the estimates obtained by regression and leads to inconsistencies in the estimates in the formation energies as they don't correspond to a thermodynamically feasible reaction system. Since these estimates, especially in the case of the group contribution method, are used for analysis of large scale metabolic networks, this inconsistency can propagate through the thermodynamic constraints imposed and lead to increased uncertainty in the thermodynamic properties of the network.

An approach to deal with the measurement error is to adjust the measurements and estimates such that the material and energy balance equations representing the system are satisfied. This is widely applied in process industries where the measurement errors can propagate through the network while modelling the process and can lead to significant errors. This approach forms the crux of data reconciliation and the estimates obtained using these methods are on an average closer to the the true values, as illustrated by the simulation studies described in the main text.

### 3 Theoretical basis of data reconciliation

The relationship between a measured value $\boldsymbol{y}$, a true value $\boldsymbol{x}$ and a random error $\boldsymbol{\varepsilon}$ can be described as [4]:

$$\boldsymbol{y} = \boldsymbol{x} + \boldsymbol{\varepsilon}$$

The random error $\boldsymbol{\varepsilon}$ in a given measurement is usually normally distributed with zero mean with variance $\boldsymbol{\sigma^2}$. The true standard deviation $\boldsymbol{\sigma}$ of the random error is not known. However, an estimate of standard deviation ($\hat{\boldsymbol{\sigma}}$ can be obtained using for the measurements when there are $\boldsymbol{N}$ samples available:

$$\hat{\boldsymbol{\sigma}} = \sqrt{\frac{1}{N-1}[\sum_{i=1}^{N}(y_i - \bar{y})^2]} \tag{5}$$

The random errors in a set of measurements can be assumed to follow a multivariate normal distribution with zero mean and a variance matrix $\boldsymbol{\Sigma}$ whose diagonal element, $\hat{\boldsymbol{\sigma}}_i^2$ is the variance of the random error in the measurement $\boldsymbol{i}$ obtained using Eq. (5). From a statistical–analysis viewpoint, the variance matrix $\boldsymbol{\Sigma}$ contains all the necessary information about the accuracy of a measurement. Thus, if the measured values are given by a vector $\boldsymbol{y}$, the most likely estimates for $\boldsymbol{x}$ can be obtained by maximizing the likelihood function of the multivariate normal distribution:

$$\max_{\mathbf{x}} \quad \frac{1}{(2\pi)^{n/2} \cdot |\Sigma|^{n/2}} \exp\left(-0.5(y-x)^T \Sigma^{-1}(y-x)\right) \tag{6}$$

where $|\boldsymbol{\Sigma}|$ is the determinant of the variance matrix. The maximum likelihood estimation problem (6) is the same as minimizing the function:

$$\min_{\mathbf{x}} \quad \sum(\mathrm{y}-\mathrm{x})\Sigma^{-1}(\mathrm{y}-\mathrm{x})^T \tag{7}$$

Often, for various physical systems, the variables are related with each other. Hence, they satisfy a set of linear or nonlinear constraints such as closure of material and energy balances or thermodynamic relationships. In data reconciliation, the information on the constraints between the true variables are also used to estimate the variables values from $\boldsymbol{y}$ such that they satisfy these constraints. Then, the objective function (Eq.(7)) is minimized subject to constraints such as material and energy balance equations, and thermodynamic constraints. This leads to a constrained least–squares optimization problem. The use of the inverse of the variance matrix in the objective function implies that the weight factor for each measurement is inversely proportional to the standard deviation of its error. This ensures that a larger weight is given to a more accurate measurement.

### 4 Construction of a projection matrix using QR decomposition

In this section, the construction of a project matrix using QR factorization $\mathbf{A_u}$ is discussed [3, 4]. The use of the QR decomposition method to identify these variables and to solve the linear data reconciliation problem (4) is briefly described here.

Consider a system with $\boldsymbol{n}$ linear constraints between $\boldsymbol{m}$ variables. The linear constraints can be written as follows

$$\mathbf{Ax{=}0}$$

where $\mathbf{A}$ is the constraint matrix and $\mathbf{0}$ is a zero vector. $\mathbf{x}$ represents the vector consisting of the reconciled estimates for the measured variables. For a partially measured system with $\boldsymbol{p}$ unmeasured variables out of the m variables, the variables can be partitioned into the measured one $\mathbf{x}_m \in \boldsymbol{R}^{m-p}$ and the unmeasured one $\mathbf{x}_u \in \boldsymbol{R}^p$ as follows:

$$\mathbf{A}_m\mathbf{x}_m + \mathbf{A}_u\mathbf{x}_u = \mathbf{0} \tag{8}$$

where $\mathbf{A}_m$ and $\mathbf{A}_u$ are of the $\boldsymbol{n} \times (\boldsymbol{m}-\boldsymbol{p})$ and $\boldsymbol{n} \times \boldsymbol{p}$-dimensional matrices, respectively. The columns of $\mathbf{A_x}$ correspond to the measured variables $\mathbf{x}_m$ while the columns of $\mathbf{A_u}$ correspond to only the unmeasured variables $\mathbf{x}_{\boldsymbol{u}}$. To obtain a reduced set of the constrained equations, a projection matrix, $\mathbf{P}$ is constructed such that $\mathbf{P}\mathbf{A}_{\boldsymbol{u}} = \mathbf{0}$. Then, the reduced set of equations from Eq. (8) can be obtained as follows:

$$\mathbf{P}\mathbf{A}_{\boldsymbol{m}}\mathbf{x}_{\boldsymbol{m}} = \mathbf{0} \tag{9}$$

Next, the projection matrix $\mathbf{P}$ will be constructed using the known matrix $\mathbf{A}_{\mathbf{u}}$. The QR decomposition of the $\mathbf{A}_{\boldsymbol{u}}$ is given by

$$\mathbf{A}_{\mathbf{u}} = \mathbf{Q}\mathbf{R} \tag{10}$$

where $\mathbf{Q}$ is the $\boldsymbol{n} \times \boldsymbol{n}$ matrix such that $\mathbf{Q}^{\boldsymbol{T}}\mathbf{Q} = \mathbf{I}$ and $\mathbf{R}$ is the $\boldsymbol{n} \times \boldsymbol{p}$-dimensional upper triangular matrix. Eq. (10) can be written as:

$$\mathbf{A}_{\boldsymbol{u}} = \mathbf{Q}\mathbf{R} = \begin{bmatrix}\mathbf{Q_1} & \mathbf{Q_2}\end{bmatrix}\begin{bmatrix}\mathbf{R_1} \\ \mathbf{0}\end{bmatrix}$$

Here, $\mathbf{R_1}$ is a $\boldsymbol{p} \times \boldsymbol{p}$-dimensional non-singular upper triangular matrix where $p$ is the rank of $\mathbf{A}_{\mathbf{u}}$. $\mathbf{R_1}$ represents the $p$ columns of $\mathbf{A}_{\mathbf{u}}$ in terms of the first $p$ basis vectors, $\mathbf{Q_1}$. Since $\mathbf{Q}$ is orthonormal, $\mathbf{Q_2^{\boldsymbol{T}}}$ has the following property:

$$\mathbf{Q_2^{\boldsymbol{T}}}\mathbf{A}_{\mathbf{u}} = \mathbf{Q_2^{\boldsymbol{T}}}\begin{bmatrix}\mathbf{Q_1} & \mathbf{Q_2}\end{bmatrix}\begin{bmatrix}\mathbf{R_1} \\ \mathbf{0}\end{bmatrix} = \begin{bmatrix}\mathbf{0} & \mathbf{I}\end{bmatrix}\begin{bmatrix}\mathbf{R_1} \\ \mathbf{0}\end{bmatrix} = \mathbf{0}$$

Thus, the projection matrix, $\mathbf{P}$ can be taken as the transpose of $\mathbf{Q_2}$. Hence, the construction of $\mathbf{P}$ can be summarized as follows: (i) Applying the QR decomposition to $\mathbf{A}_{\boldsymbol{u}}$ to compute $\mathbf{Q}$, (ii) then, define $\mathbf{Q_2}$ as the last $(\boldsymbol{n} - \boldsymbol{p})$ columns of $\mathbf{Q}$, and (iii) $\mathbf{P} = \mathbf{Q_2^{\boldsymbol{T}}}$.

### 5 Observability and Redundancy

The details related to observability and redundancy are provided in this section. Formally, the observability and redundancy are defined as follows [4].

**Definition 1** *(Observability) A variable is said to be observable if it can be estimated uniquely using the measured variables and the constraints.*

**Definition 2** *(Redundancy) A measured variable is said to be redundant if it is observable when its measurement is removed.*

A measured variable is always observable. However, the unmeasured variable is observable only if it can be estimated from the measured variables and the constraints. On the other hand, a redundant variable is always useful in improving the accuracy of the reconciled data as they can be indirectly estimated from the other measured variables and the constraints. Note that the observability and redundancy of the variables depend on the measurements and the constraints. Observability and redundancy analysis can be used for design of experiments such as choice of the set of variables to be measured for estimating a new Gibbs free energy.

The observability and redundancy analysis is performed as a part of data reconciliation framework. For a given set of variables, both measured and unmeasured, it is important to know that the observable and redundant variables. The knowledge of observable variables is useful in estimating the unmeasured variables. The observability of a variable can be easily determined from the construction of the projection matrix used in the analytical solution for a linear data reconciliation problem for a partially measured reaction system.

In Section 4 of the supporting information, it is assumed that the rows of the projection matrix $\mathbf{P}$ are independent, and hence, all the unmeasured variables are observable. If only $\boldsymbol{s}$ rows of $\mathbf{P}$ are linearly independent, then the QR decomposition of $\mathbf{A}_{\boldsymbol{u}}$ can be written as:

$$\mathbf{A}_{\boldsymbol{u}} = \mathbf{Q}\mathbf{R} = \begin{bmatrix}\mathbf{Q_1} & \mathbf{Q_2}\end{bmatrix}\begin{bmatrix}\mathbf{R_1} & \mathbf{R_2} \\ \mathbf{0} & \mathbf{0}\end{bmatrix}\mathbf{\Pi}_{\boldsymbol{u}}$$

where $\mathbf{\Pi_u}$ is a $\boldsymbol{p \times p}$-dimensional permutation matrix, i.e. the columns of $\mathbf{\Pi_u}$ consist of of the permuted columns of the identity matrix, $\mathbf{R_1}$ and $\mathbf{R_2}$ are $\boldsymbol{s \times s}$- and $\boldsymbol{s \times p - s}$-dimensional matrices, respectively. Similarly, the unmeasured variables $\mathbf{x_u}$ can be partitioned into two sets of variables:

$$\mathbf{\Pi_u x_u} = \begin{bmatrix} \mathbf{x}_{u,s} \\ \mathbf{x}_{u,p-s} \end{bmatrix}$$

By substituting the QR decomposition of $\mathbf{A_u}$ and $\mathbf{\Pi_u x_u}$ in Eq. (8), we get:

$$\mathbf{A_m x_m} + \mathbf{QR\Pi_u x_u} = \mathbf{0} \tag{11}$$

$$\mathbf{A_m x_m} + \begin{bmatrix} \mathbf{Q_1} & \mathbf{Q_2} \end{bmatrix} \begin{bmatrix} \mathbf{R_1} & \mathbf{R_2} \\ \mathbf{0} & \mathbf{0} \end{bmatrix} \begin{bmatrix} \mathbf{x}_{u,s} \\ \mathbf{x}_{u,p-s} \end{bmatrix} = \mathbf{0} \tag{12}$$

By pre-multiplying by $\mathbf{Q^T}$ to Eq. (11), the following equation can be obtained :

$$\begin{bmatrix} \mathbf{R_1} & \mathbf{R_2} \\ \mathbf{0} & \mathbf{0} \end{bmatrix} \begin{bmatrix} \mathbf{x}_{u,s} \\ \mathbf{x}_{u,p-s} \end{bmatrix} = - \begin{bmatrix} \mathbf{Q_1^T} \\ \mathbf{Q_2^T} \end{bmatrix} \mathbf{A_m x_m} \tag{13}$$

Eq. 13 can be written as follows:

$$\mathbf{x}_{u,s} = -\mathbf{R_1^{-1} Q_1^T A_m \hat{x}_m} - \mathbf{C x}_{u,p-s} \tag{14}$$

$$\mathbf{0} = \mathbf{\tilde{A}_P x_m} \tag{15}$$

where $\mathbf{C} = \mathbf{R_1^{-1} R_2}$ and $\mathbf{\tilde{A}_P} = \mathbf{Q_2^T A_m}$.

### 6 Analysis of NIST-TECR Database for Observability and Redundancy

We performed observability analysis for the entire set of reactions on the NIST-TECR database of nearly 4000 reactions between 566 compounds. It was found that the formation energy estimates of 77 compounds can be obtained entirely from the available experimental data on formation energies (of 117 compounds obtained from [2]) and the reaction equilibrium constants for 134 reactions. The data on the reaction Gibbs energies of the 134 reactions between these compounds were obtained from the NIST-TECR database. The unique estimates of these formation energies were obtained and are given in Table 1. The reconciled estimates of all 194 formation energies and 134 reactions are thermodynamically consistent, as shown in Figure 1 when measured against estimates from the linear regression.

Table 1: The Gibbs free energies of formation for 77 observable compounds using the data reconciliation approach.

| S.No | Compound | $\Delta_f G^\circ_{recon}$ (kJ/mol) | $\Delta_f G^\circ_{reg}$ (kJ/mol) |
|---|---|---|---|
| 1 | ITP | -3102.96 | -3091.00 |
| 2 | beta-Alanine | -367.72 | -373.05 |
| 3 | IDP | -2223.16 | -2223.37 |
| 4 | 2-Oxobutanoate | -459.76 | -463.81 |
| 5 | IMP | -1281.47 | -1275.87 |
| 6 | D-Alanine | -366.82 | -366.24 |
| 7 | Phenylpyruvate | -304.93 | -304.39 |
| 8 | Hydroxypyruvate | -606.61 | -607.98 |
| 9 | Carbamoyl phosphate | -1195.66 | -1210.83 |

| | | | |
|---|---|---|---|
| 10 | Cellobiose | -1582.20 | -1579.40 |
| 11 | 2-Dehydro-3-deoxy-D-gluconate | -911.39 | -914.04 |
| 12 | D-Glutamate | -693.59 | -694.95 |
| 13 | 3-Oxopropanoate | -467.08 | -471.13 |
| 14 | D-Xylulose 5-phosphate | -1591.76 | -1597.15 |
| 15 | (R)-Lactate | -514.77 | -508.59 |
| 16 | L-Homoserine | -502.30 | -490.46 |
| 17 | 5-Dehydro-D-fructose | -860.31 | -868.21 |
| 18 | D-Erythrose 4-phosphate | -1434.27 | -1435.81 |
| 19 | Orotate | -556.78 | -571.27 |
| 20 | L-Xylulose | -743.42 | -744.84 |
| 21 | (S)-Dihydroorotate | -595.14 | -606.34 |
| 22 | 6-Phospho-D-gluconate | -1957.61 | -1958.90 |
| 23 | D-Glucosamine 6-phosphate | -1594.61 | -1608.40 |
| 24 | trans-Cinnamate | -145.23 | -132.80 |
| 25 | N-Carbamoyl-L-aspartate | -783.50 | -798.58 |
| 26 | L-Aspartate 4-semialdehyde | -451.20 | -445.94 |
| 27 | Ribitol | -783.44 | -780.34 |
| 28 | (R)-Malate | -838.00 | -841.88 |
| 29 | ADP-glucose | -2597.52 | -2603.24 |
| 30 | Allantoate | -652.84 | -648.74 |
| 31 | Guanidinoacetate | -314.03 | -338.22 |
| 32 | (-)-Ureidoglycolate | -677.65 | -677.43 |
| 33 | alpha,alpha'-Trehalose 6-phosphate | -2436.37 | -2435.53 |
| 34 | 2-Oxoglutaramate | -647.23 | -642.43 |
| 35 | O-Acetyl-L-serine | -669.75 | -656.59 |
| 36 | (R)-2-Hydroxyglutarate | -844.96 | -841.23 |
| 37 | (R)-3-Hydroxybutanoate | -527.84 | -519.13 |
| 38 | D-Fructose 1-phosphate | -1753.88 | -1759.82 |
| 39 | L-Ribulose 5-phosphate | -1591.27 | -1596.66 |
| 40 | Orotidine 5'-phosphate | -1870.52 | -1882.82 |
| 41 | Trimethylamine N-oxide | 102.20 | 102.20 |
| 42 | 2-Hydroxy-3-oxopropanoate | -604.12 | -609.21 |
| 43 | 3-(4-Hydroxyphenyl)pyruvate | -468.01 | -467.48 |
| 44 | alpha-D-Glucose 1,6-bisphosphate | -2611.06 | -2605.19 |
| 45 | 2-Dehydro-3-deoxy-D-galactonate 6-phosphate | -1753.78 | -1753.93 |
| 46 | Carbamate | -348.73 | -364.53 |
| 47 | D-Leucine | -350.66 | -351.29 |
| 48 | Mesaconate | -600.96 | -600.18 |
| 49 | D-Arabitol | -782.06 | -778.96 |
| 50 | Laminaribiose | -1581.14 | -1578.34 |
| 51 | 2-Methylserine | -514.75 | -507.84 |
| 52 | 5-Oxo-D-proline | -474.85 | -472.33 |
| 53 | Phosphocreatine | -1105.83 | -1105.20 |
| 54 | (S)-2-Methylmalate | -830.83 | -833.08 |
| 55 | 2-Hydroxy-3-phenylpropenoate | -299.22 | -298.69 |
| 56 | 3-Phosphonopyruvate | -1326.92 | -1322.04 |
| 57 | 4-Phospho-L-aspartate | -1533.08 | -1536.08 |
| 58 | Phosphoguanidinoacetate | -1177.82 | -1201.38 |

| | | | |
|---|---|---|---|
| 59 | Alanopine | -665.73 | -654.81 |
| 60 | Erythrulose 1-phosphate | -1443.85 | -1437.43 |
| 61 | Adenosine tetraphosphate | -3660.37 | -3664.15 |
| 62 | Indoleglycerol phosphate | -1081.21 | -1081.02 |
| 63 | L-threo-3-Methylaspartate | -686.10 | -686.11 |
| 64 | L-5-Carboxymethylhydantoin | -597.33 | -608.54 |
| 65 | N6-(1,2-Dicarboxyethyl)-AMP | -1652.98 | -1647.85 |
| 66 | 2-Hydroxyethylenedicarboxylate | -789.21 | -791.21 |
| 67 | 2-Dehydro-3-deoxy-6-phospho-D-gluconate | -1756.20 | -1756.34 |
| 68 | Sedoheptulose 7-phosphate | -1908.22 | -1912.19 |
| 69 | Melibiose | -1577.72 | -1573.47 |
| 70 | D-arabino-Hex-3-ulose 6-phosphate | -1741.25 | -1742.09 |
| 71 | alpha-D-Glucosamine 1-phosphate | -1592.06 | -1605.86 |
| 72 | Cellotriose | -2253.56 | -2246.53 |
| 73 | 2-Oxopentanoic acid | -457.79 | -457.26 |
| 74 | 2-Keto-D-gluconic acid | -1083.20 | -1087.74 |
| 75 | Lactulose | -1575.15 | -1572.82 |
| 76 | Gentiobiose | -1585.66 | -1584.65 |
| 77 | Sucrose 6-phosphate | -2417.58 | -2419.95 |

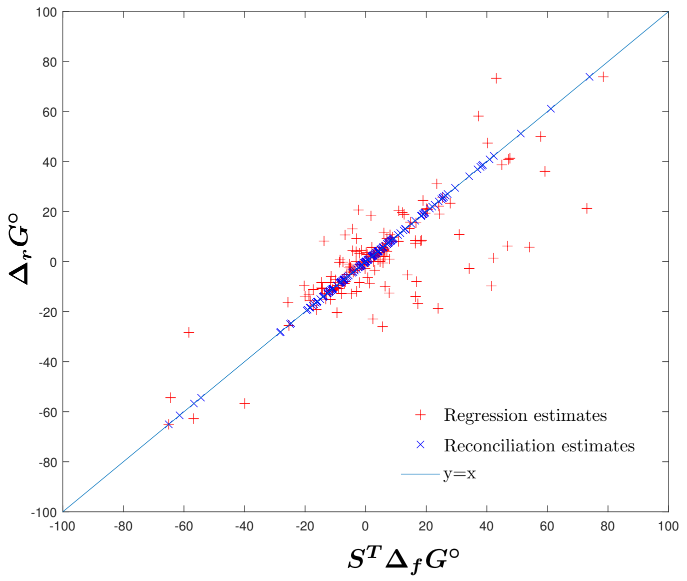

Figure 1: Comparison of the Regression and reconciliation estimates for the 194 formation energies, for which measurements are available for 117 compounds from the Alberty table. The reaction set considered is a subset of 134 reactions between these compounds obtained from the NIST-TECR database. The constraint RMSE value for the regression model for these estimates is 5.06.

### 7 Estimation of group contributions when $\mathcal{G}$ is rank deficient

The group incidence matrix for $m$ compounds given by $\mathcal{G} \in \mathbb{R}^{m \times g}$ should be full rank so that the group contributions of all $g$ groups are observable and their unique estimates obtainable. However, when $\mathcal{G}$ is rank deficient with rank $s < g$, the unobservable group contributions corresponding to the linearly dependent columns of $\mathcal{G}$ are a linear sum of the observable group contributions. Thus, in the optimization formulation, the matrix $\mathcal{G}$ can be replaced by a full rank matrix $\tilde{\mathcal{G}} \in \mathbb{R}^{m \times s}$ whose columns represent the observable group contributions. The relation between $\mathcal{G}$ and $\tilde{\mathcal{G}}$ can be expressed using a projection matrix $P_g \in \mathbb{R}^{s \times g}$ such that:

$$\mathcal{G} = \tilde{\mathcal{G}} \cdot P_g \implies P_g = (\tilde{\mathcal{G}})^{+} \cdot \mathcal{G}$$

Thus all $g$ group contributions can be obtained from the reconciliation estimates given by $\Delta_{grp}G^{\circ} \in \mathbb{R}^{s \times 1}$ using:

$$\text{Group Contributions} = (P_g)^{+} \Delta_{grp}G^{\circ}$$

### 8 Calculation of errors

#### 8.1 Constraint error

For any reaction containing $m$ compounds, the value of standard Gibbs free energy of reaction, $\Delta_r G^{\circ}$, is related to the standard formation energies of the reactants and products vector $\Delta_f G^{\circ} \in \mathbb{R}^{m \times 1}$ using a stoichiometric vector $\mathrm{s} \in \mathbb{R}^{1 \times m}$ as follows

$$\Delta_r G^{\circ} = \mathrm{s}^{\mathrm{T}} \Delta_f G^{\circ}$$

If the above equation is violated, the point on the graph plotted between $\Delta_r G^{\circ}$ and $\mathrm{s}^{T} \Delta_f G^{\circ}$ deviates from the $y = x$ line and the extent of this deviation is given by the horizontal distance of the point from this line. In this work, this is referred to as the constraint error since it quantifies the thermodynamic inconsistency of a reaction. When data reconciliation is used, the $\Delta_r G^{\circ}$ and $\Delta_f G^{\circ}$ values for the same reaction are treated as separate variables and estimates are obtained for both sets of variables. Thus, for a reconciliation estimate, the constraint error as shown in the graph is computed as the difference between $\Delta_r G^{\circ}_{est}$ and $S^T \Delta_f G^{\circ}_{est}$. For a set of $n$ reactions, the average constraint error per reaction in the reconciled estimates can be computed from:

$$\text{Constraint Error for reconciliation} = \sqrt{\frac{\sum_{i=1}^{n} \left(\Delta_r G^{\circ}_{est,i} - S_i^T \Delta_f G^{\circ}_{est,i}\right)^2}{n}}$$

However, in a linear regression model, the measured Gibbs energies of a reaction set are used to estimate all the formation energies. Since only the formation energies are estimated, the constraint error for a regression estimate is given by the residual of the fit: $\left|\Delta_r G^{\circ}_{obs} - S^T \cdot \Delta_f G^{\circ}_{est}\right|$ (see Figure 2). Thus, the average constraint error per reaction for a set of $n$ reactions for the regression method is obtained as:

$$\text{Constraint Error for regression} = \sqrt{\frac{\sum_{i=1}^{n} \left(\Delta_r G^{\circ}_{obs,i} - S_i^T \Delta_f G^{\circ}_{est,i}\right)^2}{n}}$$

#### 8.2 Prediction error

The prediction errors are quantified by the deviation of an estimate of from the true value. In this work, the simulation studied were performed by assuming the reconciled values of Gibbs free energies measurements to be the "true" values and adding the Gaussian noise to simulate experimental data. The prediction error of a reaction estimate is quantified as $\left|\Delta_r G^{\circ}_{true} - S^T \Delta_f G^{\circ}_{est}\right|$, where $\Delta_r G^{\circ}_{true}$ is the simulated "true" reaction

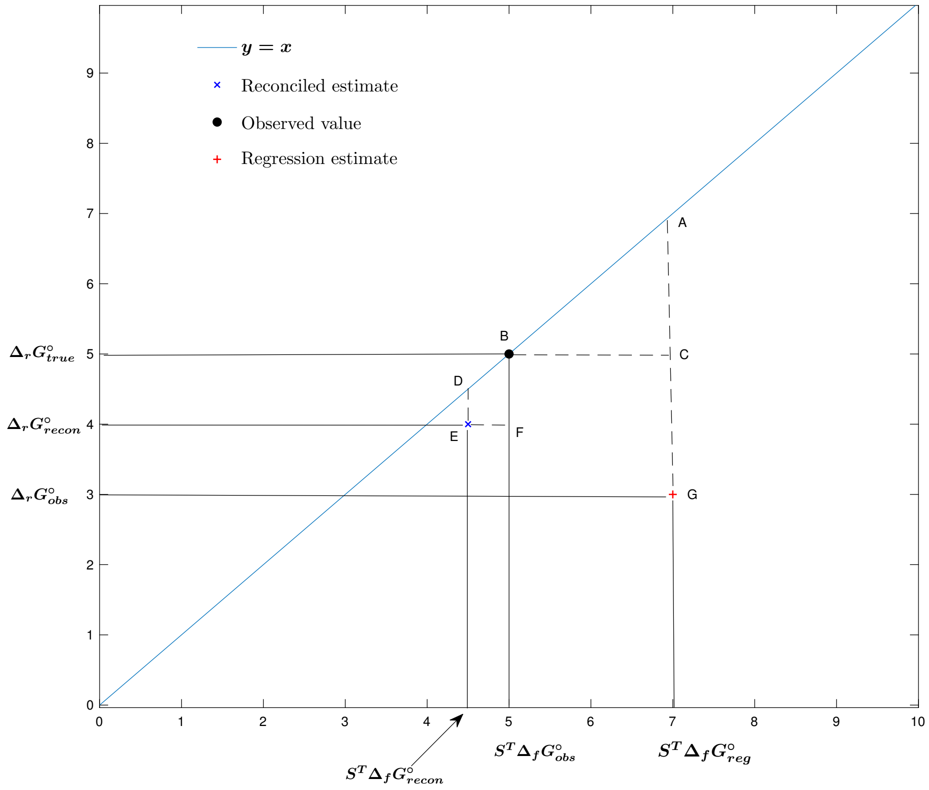

Figure 2: Graphical representation of the constraint and prediction errors. Here, the constraint error for regression estimates is given by AG which is numerically equal to $\left|\boldsymbol{\Delta_r G^\circ_{obs} - S^T \Delta_f G^\circ_{reg}}\right|$ where $\boldsymbol{\Delta_f G^\circ_{reg}}$ is the vector representing the regression estimates of the formation energies for the compounds in that reaction. The prediction error of the regression method is given by AC, which is numerically equal to $\left|\boldsymbol{\Delta_r G^\circ_{true} - S^T \Delta_f G^\circ_{reg}}\right|$. The constraint error for reconciliation estimates is given by DE and the prediction error by BF. $\boldsymbol{\Delta_r G^\circ_{recon}}$ and $\boldsymbol{\Delta_f G^\circ_{recon}}$ represent the estimates obtained by reconciliation

energy and the $\boldsymbol{\Delta_f G^\circ_{est}}$ is the estimated formation energies obtained using either the data reconciliation or regression on the simulated measurements. The average prediction error per reaction is given by:

$$\text{Prediction Error} = \sqrt{\frac{\sum_{i=1}^{n} \left(\boldsymbol{\Delta_r G^\circ_{true,i} - S_i^T \Delta_f G^\circ_{est,i}}\right)^2}{\boldsymbol{n}}}$$

For assessing the errors in the group contribution estimates, the $\boldsymbol{\Delta_f G^\circ_{est}}$ values in the above equations are replaced by $\boldsymbol{\mathcal{G} \Delta_{grp} G^\circ}$ where $\boldsymbol{\mathcal{G}}$ is the group decomposition matrix and $\boldsymbol{\Delta_{grp} G^\circ}$ is a vector representing the contribution of each group to the overall formation energy.

### 9 Experimental Data

#### 9.1 Reaction Set

The reaction network used in this work is a subset of the reactions on the NIST-DECR database, the corresponding $\boldsymbol{\Delta_r G^\circ_m}$ value for each reaction is the average of the inverse Legendre transformed apparent reaction energies of the same reaction under different conditions. If the transform is assumed to be accurate, the corresponding variance represents the accuracy of the measurement. When repeated measurements are unavailable, the variance obtained is zero and hence a unit weight is assigned to the variable when estimating its reconciled value. The reconciled estimates are represented by $\boldsymbol{\Delta_r G^\circ_{rec}}$. These values are the "true" values used in the subsequent generation of simulated measurements. Here, the regression reaction Gibbs free energy estimates, $\boldsymbol{\Delta_r G^\circ_{reg}}$, are computed as $\boldsymbol{S^T \cdot \Delta_f G^\circ_{reg}}$, where $\boldsymbol{\Delta_f G^\circ_{reg}}$ represents the formation Gibbs free energies obtained using regression (provided in Section 9.2).

Table 2: Experimental data for Reaction set obtained from the NIST-DECR database along with the reconciled estimates of Gibbs free energy of reactions $\boldsymbol{\Delta_r G^\circ_{rec}}$ and the estimates obtained by the regression approach $\boldsymbol{\Delta_r G^\circ_{reg}}$

| S.No | Reaction | $\boldsymbol{\Delta_r G^\circ_m}$ (kJ/mol) | Variance (kJ/mol) | $\boldsymbol{\Delta_r G^\circ_{rec}}$ (kJ/mol) | $\boldsymbol{\Delta_r G^\circ_{reg}}$ (kJ/mol) |
|---|---|---|---|---|---|
| 1 | isocitrate(aq) = citrate(aq) | -7.95 | 2.69 | -7.19 | -6.61 |
| 2 | ATP(aq) + D-glucose(aq) = ADP(aq) + D-glucose 6-phosphate(aq) | -13.97 | 1.62 | -10.23 | -8.03 |
| 3 | AMP(aq) + H2O(l) = adenosine(aq) + orthophosphate(aq) | -12.96 | 0.00 | -13.52 | -8.50 |
| 4 | pyrophosphate(aq) + L-serine(aq) = orthophosphate(aq) + O-phospho-L-serine(aq) | 23.36 | 0.33 | 31.11 | 26.19 |
| 5 | ATP(aq) + D-fructose 6-phosphate(aq) = ADP(aq) + D-fructose 1,6-bisphosphate(aq) | -18.68 | 7.65 | -6.76 | 3.47 |
| 6 | D-fructose 1,6-bisphosphate(aq) = 2 glycerone phosphate(aq) | 9.76 | 11.72 | 9.62 | 9.70 |
| 7 | D-glucose 6-phosphate(aq) = D-fructose 6-phosphate(aq) | 3.07 | 0.48 | 7.77 | 5.63 |
| 8 | 2 ADP(aq) = AMP(aq) + ATP(aq) | 1.44 | 1.87 | 31.63 | 23.67 |
| 9 | inosine(aq) + orthophosphate(aq) = hypoxanthine(aq) + D-ribose 1-phosphate(aq) | 9.22 | 1.12 | 10.20 | 8.29 |
| 10 | D-fructose 1,6-bisphosphate(aq) + H2O(l) = D-fructose 6-phosphate(aq) + orthophosphate(aq) | -15.62 | 28.71 | -13.81 | -22.61 |
| 11 | ATP(aq) + 3-phospho-D-glycerate(aq) = ADP(aq) + 3-phospho-D-glyceroyl phosphate(aq) | 20.34 | 1.50 | 12.39 | 11.64 |
| 12 | D-glyceraldehyde 3-phosphate(aq) = glycerone phosphate(aq) | -7.88 | 0.30 | -7.00 | -7.60 |
| 13 | ATP(aq) + pyruvate(aq) + orthophosphate(aq) = AMP(aq) + phosphoenolpyruvate(aq) + pyrophosphate(aq) | 13.67 | 37.99 | 13.19 | 14.19 |
| 14 | D-xylose(aq) = D-xylulose(aq) | 3.68 | 0.48 | 4.03 | 3.92 |
| 15 | ATP(aq) + L-glutamate(aq) + ammonia(aq) = ADP(aq) + phosphate(aq) + L-glutamine(aq) | -16.80 | 5.68 | -15.05 | -7.80 |
| 16 | D-mannose 6-phosphate(aq) = D-fructose 6-phosphate(aq) | -0.51 | 0.78 | 2.48 | 0.52 |
| 17 | ATP(aq) = adenosine 3':5'-(cyclic)phosphate(aq) + diphosphate(aq) | 6.26 | 4.30 | 7.32 | 10.53 |

| | | | | | |
|---|---|---|---|---|---|
| 18 | sucrose(aq) + orthophosphate(aq) =-D-glucose 1-phosphate(aq) + D-fructose(aq) | -8.30 | 0.25 | -9.96 | -8.96 |
| 19 | L-alanine(aq) + 2-oxoglutarate(aq) = pyruvate(aq) + L-glutamate(aq) | -0.95 | 0.98 | 2.54 | 3.15 |
| 20 | diphosphate(aq) + oxaloacetate(aq) + H2O(l) = phosphate(aq) + phosphoenolpyruvate(aq) + carbon dioxide(aq) | 50.02 | 0.00 | 43.32 | 39.28 |
| 21 | D-mannose(aq) = D-fructose(aq) | -2.50 | 0.15 | -2.13 | -3.29 |
| 22 | isocitrate(aq) = succinate(aq) + glyoxylate(aq) | 9.23 | 18.44 | 8.67 | 2.43 |
| 23 | D-galactose 1-phosphate(aq) =-D-glucose 1-phosphate(aq) | -2.79 | 0.00 | -1.95 | -1.97 |
| 24 | fumarate(aq) + H2O(l) = (S)-malate(aq) | -3.58 | 0.26 | -15.13 | -8.97 |
| 25 | D-glucose(aq) = D-fructose(aq) | -0.18 | 0.11 | 0.28 | -0.65 |
| 26 | 2-phospho-D-glycerate(aq) = 3-phospho-D-glycerate(aq) | -5.17 | 1.59 | -7.63 | -6.36 |
| 27 | D-fructose 6-phosphate(aq) + H2O(l) = D-fructose(aq) + orthophosphate(aq) | -12.81 | 1.17 | -17.83 | -17.40 |
| 28 | 3-phosphonooxypyruvate(aq) + L-glutamate(aq) = 2-oxoglutarate(aq) + O-phospho-L-serine(aq) | -12.77 | 5.12 | -12.64 | -12.12 |
| 29 | adenosine(aq) + H2O(l) = adenine(aq) + D-ribose(aq) | -9.83 | 0.00 | -13.23 | -12.94 |
| 30 | L-aspartate(aq) + 2-oxoglutarate(aq) = oxaloacetate(aq) + L-glutamate(aq) | 4.53 | 0.37 | -1.19 | -0.00 |
| 31 | -D-glucose 1-phosphate(aq) =-D-glucose 6-phosphate(aq) | -6.46 | 5.35 | -6.70 | -7.15 |
| 32 | ATP(aq) + D-ribose 5-phosphate(aq) = AMP(aq) + 5-phospho–D-ribose 1-diphosphate(aq) | -9.71 | 2.30 | -21.63 | -20.64 |
| 33 | L-aspartate(aq) = fumarate(aq) + ammonia(aq) | 15.81 | 1.66 | 14.92 | 14.11 |
| 34 | AMP(aq) + H2O(l) = adenine(aq) + D-ribose 5-phosphate(aq) | -12.86 | 0.00 | -23.86 | -15.29 |
| 35 | L-leucine(aq) + 2-oxoglutarate(aq) = 2-oxoisocaproate(aq) + L-glutamate(aq) | -3.41 | 0.45 | 0.61 | -1.20 |
| 36 | (R)-2-methylmalate(aq) = 2-methylmaleate(aq) + H2O(l) | 11.52 | 0.00 | 13.53 | 12.75 |
| 37 | D-fructose 1,6-bisphosphate(aq) = glycerone phosphate(aq) + D-glyceraldehyde 3-phosphate(aq) | 15.51 | 6.50 | 16.61 | 17.30 |
| 38 | 4-hydroxy-2-oxoglutarate(aq) = pyruvate(aq) + glyoxylate(aq) | 10.85 | 2.66 | 17.87 | 25.46 |
| 39 | lactose(aq) + H2O(l) = D-galactose(aq) + D-glucose(aq) | -9.65 | 2.92 | -11.85 | -13.39 |
| 40 | D-ribose 1-phosphate(aq) = D-ribose 5-phosphate(aq) | -8.42 | 0.00 | -17.61 | -12.19 |
| 41 | citrate(aq) = cis-aconitate(aq) + H2O(l) | 7.89 | 0.31 | 20.10 | 11.74 |
| 42 | D-ribose 5-phosphate(aq) = D-ribulose 5-phosphate(aq) | 2.83 | 1.31 | 4.29 | 3.66 |
| 43 | D-glucose 6-phosphate(aq) + H2O(l) = D-glucose(aq) + orthophosphate(aq) | -11.14 | 1.44 | -10.34 | -11.11 |
| 44 | citrate(aq) = acetate(aq) + oxaloacetate(aq) | 3.60 | 12.19 | 2.94 | -0.61 |
| 45 | alpha,alpha-Trehalose + H2O(l) = 2 D-glucose(aq) | -11.85 | 0.10 | -11.84 | -11.57 |
| 46 | isocitrate(aq) = cis-aconitate(aq) + H2O(l) | 1.44 | 0.21 | 12.91 | 5.13 |

| | | | | | |
|---|---|---|---|---|---|
| 47 | ATP(aq) + pyruvate(aq) + carbon dioxide(aq) = ADP(aq) + phosphate(aq) + oxaloacetate(aq) | -2.73 | 7.57 | -2.15 | -3.77 |
| 48 | 2-phospho-D-glycerate(aq) = phosphoenolpyruvate(aq) + H2O(l) | -3.31 | 0.45 | -9.25 | -9.85 |
| 49 | ATP(aq) + sulfate(aq) = adenosine 5'-phosphosulfate(aq) + pyrophosphate(aq) | 47.42 | 4.15 | 38.23 | 29.88 |
| 50 | adenine(aq) + 5-phospho–D-ribose 1-diphosphate(aq) = AMP(aq) + pyrophosphate(aq) | 20.66 | 9.92 | 16.45 | 8.40 |
| 51 | isomaltose(aq) + H2O(l) = 2 D-glucose(aq) | -8.73 | 4.41 | -7.98 | -6.47 |
| 52 | AMP(aq) + pyrophosphate(aq) = adenine(aq) + 5-phospho–D-ribose 1-diphosphate(aq) | -22.93 | 0.00 | -16.45 | -8.40 |
| 53 | 2-deoxy-D-ribose 5-phosphate(aq) = D-glyceraldehyde 3-phosphate(aq) + acetaldehyde(aq) | 20.94 | 0.52 | 18.04 | 19.96 |
| 54 | L-glutamine(aq) + H2O(l) = L-glutamate(aq) + ammonia(aq) | -15.08 | 0.01 | -5.52 | -11.34 |
| 55 | ATP(aq) + pyruvate(aq) = ADP(aq) + phosphoenolpyruvate(aq) | 21.29 | 19.88 | 21.65 | 22.58 |
| 56 | adenosine(aq) + orthophosphate(aq) = adenine(aq) + D-ribose 1-phosphate(aq) | 8.44 | 12.53 | 7.28 | 5.40 |
| 57 | D-arabinose 5-phosphate(aq) = D-ribulose 5-phosphate(aq) | 2.46 | 0.93 | 1.98 | 2.36 |
| 58 | O-phospho-L-serine(aq) + 2-oxoglutarate(aq) = 3-phosphonooxypyruvate(aq) + L-glutamate(aq) | 10.74 | 0.00 | 12.64 | 12.12 |
| 59 | D-lyxose(aq) = D-xylulose(aq) | 3.01 | 0.80 | 2.94 | 2.93 |
| 60 | L-tryptophan(aq) + H2O(l) = indole(aq) + pyruvate(aq) + ammonia(aq) | 22.22 | 1.20 | 23.02 | 24.40 |
| 61 | (S)-lactate(aq) + oxaloacetate(aq) = (S)-malate(aq) + pyruvate(aq) | -2.54 | 2.25 | -0.50 | 2.71 |
| 62 | pyruvate(aq) + orthophosphate(aq) = acetyl phosphate(aq) + formate(aq) | -8.36 | 0.00 | -7.22 | -8.39 |
| 63 | D-mannose 6-phosphate(aq) + H2O(l) = D-mannose(aq) + orthophosphate(aq) | -9.48 | 0.00 | -13.22 | -13.59 |
| 64 | D-mannose 1-phosphate(aq) = D-mannose 6-phosphate(aq) | -4.13 | 13.50 | -4.22 | -5.54 |
| 65 | ATP(aq) + acetate(aq) = ADP(aq) + acetyl phosphate(aq) | 13.29 | 2.54 | 8.94 | 7.00 |
| 66 | (R)-3-phosphoglycerate(aq) + H2O(l) = (R)-glycerate(aq) + orthophosphate(aq) | -17.69 | 0.09 | -17.09 | -18.02 |
| 67 | trehalose(aq) + orthophosphate(aq) = D-glucose(aq) + D-glucose 1-phosphate(aq) | 4.52 | 3.27 | 5.21 | 6.69 |
| 68 | glycine(aq) + formaldehyde(aq) = L-serine(aq) | -20.37 | 0.00 | -15.01 | -15.07 |
| 69 | D-ribose(aq) = D-ribulose(aq) | 1.99 | 0.12 | 1.19 | 1.97 |
| 70 | adenosine(aq) + H2O(l) = inosine(aq) + ammonia(aq) | -62.75 | 8.06 | -61.74 | -59.48 |
| 71 | pyrophosphate(aq) + H2O(l) = 2 orthophosphate(aq) | 24.47 | 15.32 | 19.53 | 12.93 |
| 72 | D-fructose(aq) + D-glyceraldehyde-3-phosphate(aq) = D-fructose 6-phosphate(aq) + D-glyceraldehyde(aq) | 3.31 | 0.00 | 3.91 | 1.74 |

| | | | | | |
|---|---|---|---|---|---|
| 73 | ATP(aq) + sulfate(aq) + H2O(l) = 2 orthophosphate(aq) + adenosine 5'-phosphosulfate(aq) | 36.09 | 0.00 | 57.76 | 42.81 |
| 74 | ATP(aq) + H2O(l) = ADP(aq) + orthophosphate(aq) | -25.99 | 0.00 | -20.57 | -19.14 |
| 75 | ATP(aq) + pyruvate(aq) + H2O(l) = AMP(aq) + phosphoenolpyruvate(aq) + orthophosphate(aq) | -10.61 | 0.00 | 32.71 | 27.12 |
| 76 | maltose(aq) + orthophosphate(aq) = D-glucose(aq) + D-glucose 1-phosphate(aq) | 3.36 | 0.00 | 1.99 | 2.96 |
| 77 | maltose(aq) + H2O(l) = 2 D-glucose(aq) | -13.78 | 0.00 | -15.06 | -15.30 |
| 78 | pyrophosphate(aq) + D-fructose 6-phosphate(aq) = orthophosphate(aq) + D-fructose 1,6-bisphosphate(aq) | 58.20 | 0.00 | 33.34 | 35.54 |
| 79 | D-galactose 6-phosphate(aq) + H2O(l) = D-galactose(aq) + orthophosphate(aq) | -11.55 | 0.00 | -11.80 | -12.52 |
| 80 | L-O-phosphoserine(aq) + H2O(l) = L-serine(aq) + orthophosphate(aq) | -10.76 | 3.99 | -11.59 | -13.26 |
| 81 | D-arabinose(aq) = D-ribulose(aq) | 5.43 | 1.61 | 5.59 | 5.55 |
| 82 | glycine(aq) + acetaldehyde(aq) = L-threonine(aq) | -10.29 | 0.00 | -10.91 | -10.62 |
| 83 | inosine(aq) + H2O(l) = hypoxanthine(aq) + D-ribose(aq) | -11.91 | 0.00 | -10.30 | -10.05 |
| 84 | D-arabinose(aq) = D-ribose(aq) | 3.06 | 0.19 | 4.41 | 3.58 |
| 85 | D-glucose(aq) = D-mannose(aq) | 2.03 | 0.00 | 2.41 | 2.63 |
| 86 | ATP(aq) + D-galactose(aq) = ADP(aq) + D-galactose 1-phosphate(aq) | -7.97 | 0.02 | -8.32 | -6.30 |
| 87 | glycine(aq) + oxaloacetate(aq) = glyoxylate(aq) + L-aspartate(aq) | 8.63 | 0.00 | 4.54 | 3.96 |

#### 9.2 Compounds

Experimental data on formation energies was obtained from [2]. As measurements of apparent formation energies of the same compound under different conditions are unavailable, the inverse Legendre transformed values are directly used and a unit weight is assigned while reconciling the measurements.

Table 3: Experimental values of compound formation energies under standard conditions obtained from [2] along with the Gibbs free energies of formations estimates obtained by applying the data reconciliation approach ($\boldsymbol{\Delta_f G^\circ_{rec}}$) and the regression method $\boldsymbol{\Delta_f G^\circ_{reg}}$.

| S.No | Compound | $\boldsymbol{\Delta_f G^\circ_m}$ | $\boldsymbol{\Delta_f G^\circ_{rec}}$ | $\boldsymbol{\Delta_f G^\circ_{reg}}$ |
|---|---|---|---|---|
| 1 | H2O | -237.19 | -225.67 | -227.99 |
| 2 | ATP | -2811.57 | -2781.15 | -2782.14 |
| 3 | ADP | -1947.09 | -1935.12 | -1933.51 |
| 4 | Orthophosphate | -1096.07 | -1092.27 | -1095.76 |
| 5 | Diphosphate | -1973.88 | -1978.39 | -1976.46 |
| 6 | NH3 | -79.33 | -77.10 | -77.83 |
| 7 | AMP | -1040.48 | -1057.45 | -1061.20 |
| 8 | Pyruvate | -472.22 | -462.14 | -459.89 |
| 9 | L-Glutamate | -697.52 | -691.98 | -694.78 |
| 10 | 2-Oxoglutarate | -793.44 | -789.10 | -790.93 |
| 11 | D-Glucose | -915.93 | -909.03 | -909.94 |
| 12 | Acetate | -369.32 | -372.36 | -371.40 |

| | | | | |
|---|---|---|---|---|
| 13 | Oxaloacetate | -793.25 | -793.73 | -793.63 |
| 14 | Glycine | -379.89 | -379.24 | -379.58 |
| 15 | L-Alanine | -370.99 | -367.56 | -366.89 |
| 16 | Succinate | -690.41 | -680.16 | -683.60 |
| 17 | Glyoxylate | -468.63 | -473.01 | -471.77 |
| 18 | L-Aspartate | -695.92 | -695.42 | -697.48 |
| 19 | Formate | -351.03 | -352.17 | -351.00 |
| 20 | Sulfate | -744.56 | -761.02 | -755.38 |
| 21 | L-Glutamine | -528.03 | -537.90 | -533.28 |
| 22 | L-Serine | -510.85 | -510.37 | -510.84 |
| 23 | Formaldehyde | -121.49 | -116.12 | -116.18 |
| 24 | Phosphoenolpyruvate | -1263.67 | -1286.52 | -1285.94 |
| 25 | L-Tryptophan | -114.71 | -113.76 | -112.53 |
| 26 | Acetaldehyde | -138.98 | -138.08 | -138.33 |
| 27 | D-Fructose 6-phosphate | -1760.82 | -1757.51 | -1760.97 |
| 28 | Sucrose | -1564.67 | -1565.09 | -1565.33 |
| 29 | D-Glucose 6-phosphate | -1763.93 | -1765.28 | -1766.60 |
| 30 | D-Fructose | -915.53 | -908.74 | -910.59 |
| 31 | D-Glucose | -1756.84 | -1758.58 | -1759.46 |
| 32 | Glycerone phosphate | -1296.27 | -1300.34 | -1298.22 |
| 33 | D-Ribose 5-phosphate | -1595.74 | -1601.00 | -1599.64 |
| 34 | D-Glyceraldehyde 3-phosphate | -1288.51 | -1293.34 | -1290.62 |
| 35 | 5-Phospho-alpha-D-ribose 1-diphosphate | -3325.36 | -3346.32 | -3341.23 |
| 36 | D-Ribose | -738.74 | -737.30 | -738.02 |
| 37 | Fumarate | -601.85 | -603.40 | -605.53 |
| 38 | L-Leucine | -352.26 | -350.47 | -350.05 |
| 39 | D-Galactose | -908.93 | -902.28 | -902.55 |
| 40 | Adenine | 313.39 | 294.02 | 295.17 |
| 41 | (S)-Malate | -842.63 | -844.20 | -842.49 |
| 42 | Citrate | -1162.73 | -1169.03 | -1164.42 |
| 43 | D-Mannose | -910.03 | -906.61 | -907.31 |
| 44 | D-Xylose | -750.44 | -750.28 | -750.21 |
| 45 | (S)-Lactate | -516.71 | -512.11 | -511.46 |
| 46 | L-Threonine | -528.85 | -528.23 | -528.53 |
| 47 | 3-Phospho-D-glycerate | -1502.59 | -1510.57 | -1510.44 |
| 48 | D-Ribulose 5-phosphate | -1595.26 | -1596.71 | -1595.98 |
| 49 | Maltose | -1574.67 | -1577.32 | -1576.59 |
| 50 | Adenosine | -194.57 | -204.38 | -201.92 |
| 51 | D-Arabinose | -742.24 | -741.71 | -741.60 |
| 52 | Adenylyl sulfate | -1541.99 | -1525.54 | -1531.18 |
| 53 | Acetyl phosphate | -1219.37 | -1209.46 | -1213.04 |
| 54 | 4-Methyl-2-oxopentanoate | -445.19 | -446.98 | -447.40 |
| 55 | 3-Phospho-D-glyceroyl phosphate | -2356.14 | -2344.21 | -2347.44 |
| 56 | Lactose | -1567.37 | -1573.79 | -1571.11 |
| 57 | Isomaltose | -1587.67 | -1584.40 | -1585.42 |
| 58 | D-Glycerate | -661.01 | -661.06 | -660.68 |
| 59 | Hypoxanthine | 89.36 | 86.65 | 88.42 |
| 60 | D-Mannose 6-phosphate | -1759.83 | -1759.99 | -1761.49 |
| 61 | HCO3 | -586.79 | -575.68 | -577.09 |

| | | | | |
|---|---|---|---|---|
| 62 | Inosine | -409.23 | -414.68 | -411.56 |
| 63 | D-Ribulose | -735.94 | -736.12 | -736.05 |
| 64 | D-Xylulose | -746.14 | -746.25 | -746.29 |
| 65 | Isocitrate | -1156.03 | -1161.84 | -1157.80 |
| 66 | D-Fructose | -2601.40 | -2610.30 | -2606.14 |
| 67 | cis-Aconitate | -917.14 | -923.26 | -924.68 |
| 68 | Indole | 223.79 | 222.83 | 221.61 |
| 69 | D-Lyxose | -749.14 | -749.20 | -749.22 |
| 70 | 3',5'-Cyclic AMP | -790.88 | -795.43 | -795.15 |
| 71 | D-Glyceraldehyde | -440.07 | -440.66 | -438.50 |
| 72 | alpha-D-Ribose 1-phosphate | -1587.65 | -1583.39 | -1587.46 |
| 73 | 2-Phospho-D-glycerate | -1496.36 | -1502.94 | -1504.08 |
| 74 | D-Mannose 1-phosphate | -1754.54 | -1755.77 | -1755.95 |
| 75 | 2-Deoxy-D-ribose 5-phosphate | -1447.94 | -1449.46 | -1448.91 |
| 76 | O-Phospho-L-serine | -1360.76 | -1365.38 | -1365.35 |
| 77 | alpha,alpha-Trehalose | -1582.77 | -1580.54 | -1580.33 |
| 78 | D-Arabinose 5-phosphate | -1598.25 | -1598.70 | -1598.35 |
| 79 | D-Galactose 6-phosphate | -1756.83 | -1757.08 | -1757.80 |
| 80 | 4-Hydroxy-2-oxoglutarate | -971.74 | -953.02 | -957.12 |
| 81 | 2-Methylmaleate | -600.71 | -602.71 | -601.94 |
| 82 | (R)-2-Methylmalate | -843.91 | -841.91 | -842.68 |
| 83 | 3-Phosphonooxypyruvate | -1448.65 | -1449.87 | -1449.38 |
| 84 | D-Galactose 1-phosphate | -1756.64 | -1756.63 | -1757.49 |

### 10 Calculation of confidence intervals using a Monte Carlo simulation for reconciled data

To determine the uncertainty of the estimates obtained by the data reconciliation, a Monte Carlo simulation was used. For each "true" Gibbs energy value, i.e. reconciled value, in the reaction set considered in this study, 100 outcomes were generated by adding Gaussian noise with $\boldsymbol{\alpha} = \mathbf{10}$. The DR framework was then applied to each of the 100 sets of simulated measurements and the variability in the resulting distribution of reconciled estimates was quantified by the standard error (SE) and confidence intervals (CI). The results of the Monte-carlo simulation are given in the following sections for the two approaches discussed in this work: (i) data reconciliation estimates for full set of measurements, and (ii) group contribution estimates.

#### 10.1 Data reconciliation estimates

##### 10.1.1 Reactions

| Confidence intervals of reaction Gibbs free energy estimates | | | | |
|---|---|---|---|---|
| S.No | Reaction | Mean | SE | CI |
| 1 | isocitrate(aq) = citrate(aq) | -7.33 | 1.29 | (-9.85,-4.81) |
| 2 | ATP(aq) + D-glucose(aq) = ADP(aq) + D-glucose 6-phosphate(aq) | -8.94 | 2.50 | (-13.84,-4.03) |
| 3 | AMP(aq) + H2O(l) = adenosine(aq) + orthophosphate(aq) | -13.75 | 1.03 | (-15.76,-11.73) |
| 4 | pyrophosphate(aq) + L-serine(aq) = orthophosphate(aq) + O-phospho-L-serine(aq) | 31.64 | 1.98 | (27.77,35.52) |

| | | | | |
|---|---|---|---|---|
| 5 | ATP(aq) + D-fructose 6-phosphate(aq) = ADP(aq) + D-fructose 1,6-bisphosphate(aq) | -3.16 | 2.87 | (-8.79,2.47) |
| 6 | D-fructose 1,6-bisphosphate(aq) = 2 glycerone phosphate(aq) | 8.45 | 2.91 | (2.75,14.14) |
| 7 | D-glucose 6-phosphate(aq) = D-fructose 6-phosphate(aq) | 6.75 | 1.92 | (2.99,10.51) |
| 8 | 2 ADP(aq) = AMP(aq) + ATP(aq) | 31.93 | 2.68 | (26.69,37.18) |
| 9 | inosine(aq) + orthophosphate(aq) = hypoxanthine(aq) + D-ribose 1-phosphate(aq) | 10.57 | 1.24 | (8.13,13.01) |
| 10 | D-fructose 1,6-bisphosphate(aq) + H2O(l) = D-fructose 6-phosphate(aq) + orthophosphate(aq) | -17.67 | 2.04 | (-21.67,-13.66) |
| 11 | ATP(aq) + 3-phospho-D-glycerate(aq) = ADP(aq) + 3-phospho-D-glyceroyl phosphate(aq) | 13.20 | 2.39 | (8.52,17.89) |
| 12 | D-glyceraldehyde 3-phosphate(aq) = glycerone phosphate(aq) | -6.54 | 1.77 | (-10.00,-3.08) |
| 13 | ATP(aq) + pyruvate(aq) + orthophosphate(aq) = AMP(aq) + phosphoenolpyruvate(aq) + pyrophosphate(aq) | 14.37 | 2.14 | (10.18,18.56) |
| 14 | D-xylose(aq) = D-xylulose(aq) | 4.66 | 1.25 | (2.21,7.12) |
| 15 | ATP(aq) + L-glutamate(aq) + ammonia(aq) = ADP(aq) + phosphate(aq) + L-glutamine(aq) | -14.40 | 1.66 | (-17.65,-11.14) |
| 16 | D-mannose 6-phosphate(aq) = D-fructose 6-phosphate(aq) | 3.29 | 2.02 | (-0.67,7.25) |
| 17 | ATP(aq) = adenosine 3':5'-(cyclic)phosphate(aq) + diphosphate(aq) | 8.77 | 2.10 | (4.64,12.89) |
| 18 | sucrose(aq) + orthophosphate(aq) =-D-glucose 1-phosphate(aq) + D-fructose(aq) | -10.15 | 2.48 | (-15.01,-5.29) |
| 19 | L-alanine(aq) + 2-oxoglutarate(aq) = pyruvate(aq) + L-glutamate(aq) | 1.65 | 1.34 | (-0.98,4.27) |
| 20 | diphosphate(aq) + oxaloacetate(aq) + H2O(l) = phosphate(aq) + phosphoenolpyruvate(aq) + carbon dioxide(aq) | 44.66 | 1.72 | (41.28,48.03) |
| 21 | D-mannose(aq) = D-fructose(aq) | -2.90 | 1.41 | (-5.65,-0.14) |
| 22 | isocitrate(aq) = succinate(aq) + glyoxylate(aq) | 10.37 | 1.46 | (7.52,13.23) |
| 23 | D-galactose 1-phosphate(aq) =-D-glucose 1-phosphate(aq) | -1.43 | 1.89 | (-5.13,2.26) |
| 24 | fumarate(aq) + H2O(l) = (S)-malate(aq) | -17.06 | 1.18 | (-19.38,-14.75) |
| 25 | D-glucose(aq) = D-fructose(aq) | 0.79 | 1.23 | (-1.62,3.21) |
| 26 | 2-phospho-D-glycerate(aq) = 3-phospho-D-glycerate(aq) | -7.44 | 1.64 | (-10.66,-4.22) |
| 27 | D-fructose 6-phosphate(aq) + H2O(l) = D-fructose(aq) + orthophosphate(aq) | -17.84 | 1.69 | (-21.14,-14.54) |
| 28 | 3-phosphonooxypyruvate(aq) + L-glutamate(aq) = 2-oxoglutarate(aq) + O-phospho-L-serine(aq) | -13.73 | 1.72 | (-17.09,-10.37) |
| 29 | adenosine(aq) + H2O(l) = adenine(aq) + D-ribose(aq) | -12.20 | 0.97 | (-14.10,-10.31) |
| 30 | L-aspartate(aq) + 2-oxoglutarate(aq) = oxaloacetate(aq) + L-glutamate(aq) | -2.84 | 1.46 | (-5.72,0.03) |
| 31 | -D-glucose 1-phosphate(aq) =-D-glucose 6-phosphate(aq) | -6.34 | 1.97 | (-10.20,-2.47) |

| | | | | |
|---|---|---|---|---|
| 32 | ATP(aq) + D-ribose 5-phosphate(aq) = AMP(aq) + 5-phospho–D-ribose 1-diphosphate(aq) | -18.46 | 2.79 | (-23.93,-12.99) |
| 33 | L-aspartate(aq) = fumarate(aq) + ammonia(aq) | 15.86 | 1.03 | (13.85,17.87) |
| 34 | AMP(aq) + H2O(l) = adenine(aq) + D-ribose 5-phosphate(aq) | -23.94 | 1.46 | (-26.80,-21.09) |
| 35 | L-leucine(aq) + 2-oxoglutarate(aq) = 2-oxoisocaproate(aq) + L-glutamate(aq) | 0.41 | 1.37 | (-2.28,3.09) |
| 36 | (R)-2-methylmalate(aq) = 2-methylmaleate(aq) + H2O(l) | 13.53 | 1.08 | (11.41,15.65) |
| 37 | D-fructose 1,6-bisphosphate(aq) = glycerone phosphate(aq) + D-glyceraldehyde 3-phosphate(aq) | 14.99 | 2.35 | (10.39,19.59) |
| 38 | 4-hydroxy-2-oxoglutarate(aq) = pyruvate(aq) + glyoxylate(aq) | 20.52 | 1.33 | (17.93,23.12) |
| 39 | lactose(aq) + H2O(l) = D-galactose(aq) + D-glucose(aq) | -12.75 | 1.78 | (-16.25,-9.25) |
| 40 | D-ribose 1-phosphate(aq) = D-ribose 5-phosphate(aq) | -18.58 | 1.54 | (-21.60,-15.56) |
| 41 | citrate(aq) = cis-aconitate(aq) + H2O(l) | 20.38 | 1.46 | (17.51,23.24) |
| 42 | D-ribose 5-phosphate(aq) = D-ribulose 5-phosphate(aq) | 3.36 | 1.65 | (0.14,6.59) |
| 43 | D-glucose 6-phosphate(aq) + H2O(l) = D-glucose(aq) + orthophosphate(aq) | -11.88 | 1.69 | (-15.19,-8.58) |
| 44 | citrate(aq) = acetate(aq) + oxaloacetate(aq) | 2.30 | 1.51 | (-0.65,5.25) |
| 45 | alpha,alpha-Trehalose + H2O(l) = 2 D-glucose(aq) | -13.57 | 2.01 | (-17.51,-9.63) |
| 46 | isocitrate(aq) = cis-aconitate(aq) + H2O(l) | 13.05 | 1.50 | (10.11,15.98) |
| 47 | ATP(aq) + pyruvate(aq) + carbon dioxide(aq) = ADP(aq) + phosphate(aq) + oxaloacetate(aq) | -3.85 | 2.15 | (-8.05,0.36) |
| 48 | 2-phospho-D-glycerate(aq) = phosphoenolpyruvate(aq) + H2O(l) | -9.56 | 1.85 | (-13.18,-5.93) |
| 49 | ATP(aq) + sulfate(aq) = adenosine 5'-phosphosulfate(aq) + pyrophosphate(aq) | 38.81 | 2.09 | (34.71,42.90) |
| 50 | adenine(aq) + 5-phospho–D-ribose 1-diphosphate(aq) = AMP(aq) + pyrophosphate(aq) | 13.91 | 2.17 | (9.66,18.15) |
| 51 | isomaltose(aq) + H2O(l) = 2 D-glucose(aq) | -9.54 | 2.05 | (-13.55,-5.52) |
| 52 | AMP(aq) + pyrophosphate(aq) = adenine(aq) + 5-phospho–D-ribose 1-diphosphate(aq) | -13.91 | 2.17 | (-18.15,-9.66) |
| 53 | 2-deoxy-D-ribose 5-phosphate(aq) = D-glyceraldehyde 3-phosphate(aq) + acetaldehyde(aq) | 20.15 | 1.49 | (17.22,23.07) |
| 54 | L-glutamine(aq) + H2O(l) = L-glutamate(aq) + ammonia(aq) | -6.43 | 0.98 | (-8.35,-4.50) |
| 55 | ATP(aq) + pyruvate(aq) = ADP(aq) + phosphoenolpyruvate(aq) | 22.04 | 1.86 | (18.40,25.67) |
| 56 | adenosine(aq) + orthophosphate(aq) = adenine(aq) + D-ribose 1-phosphate(aq) | 8.39 | 1.31 | (5.82,10.95) |
| 57 | D-arabinose 5-phosphate(aq) = D-ribulose 5-phosphate(aq) | 0.95 | 1.54 | (-2.07,3.96) |
| 58 | O-phospho-L-serine(aq) + 2-oxoglutarate(aq) = 3-phosphonooxypyruvate(aq) + L-glutamate(aq) | 13.73 | 1.72 | (10.37,17.09) |
| 59 | D-lyxose(aq) = D-xylulose(aq) | 4.57 | 1.00 | (2.61,6.54) |

| | | | | |
|---|---|---|---|---|
| 60 | L-tryptophan(aq) + H2O(l) = indole(aq) + pyruvate(aq) + ammonia(aq) | 23.86 | 0.57 | (22.74,24.98) |
| 61 | (S)-lactate(aq) + oxaloacetate(aq) = (S)-malate(aq) + pyruvate(aq) | -0.27 | 1.39 | (-2.99,2.45) |
| 62 | pyruvate(aq) + orthophosphate(aq) = acetyl phosphate(aq) + formate(aq) | -7.09 | 1.56 | (-10.15,-4.03) |
| 63 | D-mannose 6-phosphate(aq) + H2O(l) = D-mannose(aq) + orthophosphate(aq) | -11.66 | 1.70 | (-15.00,-8.32) |
| 64 | D-mannose 1-phosphate(aq) = D-mannose 6-phosphate(aq) | -6.07 | 1.78 | (-9.56,-2.59) |
| 65 | ATP(aq) + acetate(aq) = ADP(aq) + acetyl phosphate(aq) | 9.31 | 2.08 | (5.24,13.39) |
| 66 | (R)-3-phosphoglycerate(aq) + H2O(l) = (R)-glycerate(aq) + orthophosphate(aq) | -18.09 | 1.43 | (-20.89,-15.28) |
| 67 | trehalose(aq) + orthophosphate(aq) = D-glucose(aq) + D-glucose 1-phosphate(aq) | 4.65 | 2.14 | (0.45,8.85) |
| 68 | glycine(aq) + formaldehyde(aq) = L-serine(aq) | -14.03 | 0.74 | (-15.49,-12.57) |
| 69 | D-ribose(aq) = D-ribulose(aq) | 0.55 | 1.20 | (-1.81,2.91) |
| 70 | adenosine(aq) + H2O(l) = inosine(aq) + ammonia(aq) | -61.47 | 0.47 | (-62.39,-60.55) |
| 71 | pyrophosphate(aq) + H2O(l) = 2 orthophosphate(aq) | 18.78 | 1.59 | (15.67,21.88) |
| 72 | D-fructose(aq) + D-glyceraldehyde-3-phosphate(aq) = D-fructose 6-phosphate(aq) + D-glyceraldehyde(aq) | 3.09 | 2.09 | (-1.00,7.19) |
| 73 | ATP(aq) + sulfate(aq) + H2O(l) = 2 orthophosphate(aq) + adenosine 5'-phosphosulfate(aq) | 57.59 | 1.65 | (54.35,60.82) |
| 74 | ATP(aq) + H2O(l) = ADP(aq) + orthophosphate(aq) | -20.82 | 1.60 | (-23.95,-17.69) |
| 75 | ATP(aq) + pyruvate(aq) + H2O(l) = AMP(aq) + phosphoenolpyruvate(aq) + orthophosphate(aq) | 33.15 | 1.48 | (30.24,36.05) |
| 76 | maltose(aq) + orthophosphate(aq) = D-glucose(aq) + D-glucose 1-phosphate(aq) | 2.44 | 2.04 | (-1.56,6.44) |
| 77 | maltose(aq) + H2O(l) = 2 D-glucose(aq) | -15.78 | 1.88 | (-19.46,-12.10) |
| 78 | pyrophosphate(aq) + D-fructose 6-phosphate(aq) = orthophosphate(aq) + D-fructose 1,6-bisphosphate(aq) | 36.44 | 2.30 | (31.94,40.95) |
| 79 | D-galactose 6-phosphate(aq) + H2O(l) = D-galactose(aq) + orthophosphate(aq) | -14.38 | 1.69 | (-17.70,-11.07) |
| 80 | L-O-phosphoserine(aq) + H2O(l) = L-serine(aq) + orthophosphate(aq) | -12.86 | 1.41 | (-15.63,-10.10) |
| 81 | D-arabinose(aq) = D-ribulose(aq) | 5.85 | 1.22 | (3.46,8.24) |
| 82 | glycine(aq) + acetaldehyde(aq) = L-threonine(aq) | -11.12 | 1.00 | (-13.09,-9.16) |
| 83 | inosine(aq) + H2O(l) = hypoxanthine(aq) + D-ribose(aq) | -10.02 | 0.99 | (-11.96,-8.08) |
| 84 | D-arabinose(aq) = D-ribose(aq) | 5.30 | 1.12 | (3.10,7.49) |
| 85 | D-glucose(aq) = D-mannose(aq) | 3.69 | 1.26 | (1.21,6.16) |
| 86 | ATP(aq) + D-galactose(aq) = ADP(aq) + D-galactose 1-phosphate(aq) | -7.80 | 2.18 | (-12.07,-3.52) |

| | | | | |
|---|---|---|---|---|
| 87 | glycine(aq) + oxaloacetate(aq) = glyoxylate(aq) + L-aspartate(aq) | 7.24 | 1.38 | (4.54,9.93) |

##### 10.1.2 Compounds

| Confidence intervals of formation Gibbs free energy estimates | | | | |
|---|---|---|---|---|
| S.No | Compound | Mean | SE | CI |
| 1 | H2O | -225.57 | 0.52 | (-226.60,-224.55) |
| 2 | ATP | -2782.97 | 1.53 | (-2785.96,-2779.98) |
| 3 | ADP | -1936.45 | 1.21 | (-1938.81,-1934.08) |
| 4 | Orthophosphate | -1092.92 | 0.80 | (-1094.50,-1091.35) |
| 5 | Diphosphate | -1979.05 | 1.31 | (-1981.63,-1976.47) |
| 6 | NH3 | -76.94 | 0.43 | (-77.78,-76.11) |
| 7 | AMP | -1057.99 | 0.84 | (-1059.62,-1056.35) |
| 8 | Pyruvate | -461.87 | 0.58 | (-463.01,-460.73) |
| 9 | L-Glutamate | -692.84 | 0.83 | (-694.47,-691.22) |
| 10 | 2-Oxoglutarate | -789.35 | 0.88 | (-791.08,-787.62) |
| 11 | D-Glucose | -909.67 | 0.80 | (-911.23,-908.11) |
| 12 | Acetate | -372.62 | 0.59 | (-373.77,-371.47) |
| 13 | Oxaloacetate | -794.69 | 0.76 | (-796.19,-793.19) |
| 14 | Glycine | -379.62 | 0.50 | (-380.60,-378.64) |
| 15 | L-Alanine | -367.01 | 0.67 | (-368.31,-365.70) |
| 16 | Succinate | -680.17 | 0.82 | (-681.77,-678.57) |
| 17 | Glyoxylate | -471.74 | 0.61 | (-472.93,-470.55) |
| 18 | L-Aspartate | -695.34 | 0.74 | (-696.78,-693.89) |
| 19 | Formate | -352.05 | 0.58 | (-353.18,-350.92) |
| 20 | Sulfate | -760.65 | 0.83 | (-762.28,-759.01) |
| 21 | L-Glutamine | -537.78 | 0.77 | (-539.29,-536.28) |
| 22 | L-Serine | -510.16 | 0.62 | (-511.37,-508.95) |
| 23 | Formaldehyde | -116.51 | 0.31 | (-117.10,-115.91) |
| 24 | Phosphoenolpyruvate | -1286.36 | 1.20 | (-1288.70,-1284.01) |
| 25 | L-Tryptophan | -113.86 | 0.27 | (-114.40,-113.33) |
| 26 | Acetaldehyde | -137.63 | 0.35 | (-138.31,-136.94) |
| 27 | D-Fructose 6-phosphate | -1758.39 | 1.36 | (-1761.05,-1755.73) |
| 28 | Sucrose | -1564.61 | 1.35 | (-1567.25,-1561.96) |
| 29 | D-Glucose 6-phosphate | -1765.14 | 1.43 | (-1767.94,-1762.33) |
| 30 | D-Fructose | -908.88 | 0.94 | (-910.72,-907.04) |
| 31 | D-Glucose | -1758.80 | 1.26 | (-1761.28,-1756.33) |
| 32 | Glycerone phosphate | -1299.81 | 1.20 | (-1302.16,-1297.47) |
| 33 | D-Ribose 5-phosphate | -1601.70 | 1.15 | (-1603.96,-1599.44) |
| 34 | D-Glyceraldehyde 3-phosphate | -1293.27 | 1.15 | (-1295.51,-1291.03) |
| 35 | 5-Phospho-alpha-D-ribose 1-diphosphate | -3345.14 | 1.72 | (-3348.51,-3341.78) |
| 36 | D-Ribose | -736.36 | 0.77 | (-737.86,-734.86) |
| 37 | Fumarate | -602.53 | 0.82 | (-604.14,-600.93) |
| 38 | L-Leucine | -350.99 | 0.56 | (-352.08,-349.89) |
| 39 | D-Galactose | -903.05 | 0.87 | (-904.74,-901.35) |
| 40 | Adenine | 294.19 | 0.45 | (293.31,295.08) |
| 41 | (S)-Malate | -845.17 | 0.93 | (-846.99,-843.36) |

| | | | | |
|---|---|---|---|---|
| 42 | Citrate | -1169.61 | 0.93 | (-1171.43,-1167.79) |
| 43 | D-Mannose | -905.99 | 0.96 | (-907.87,-904.10) |
| 44 | D-Xylose | -749.70 | 0.74 | (-751.15,-748.24) |
| 45 | (S)-Lactate | -512.08 | 0.69 | (-513.43,-510.73) |
| 46 | L-Threonine | -528.37 | 0.73 | (-529.79,-526.95) |
| 47 | 3-Phospho-D-glycerate | -1509.82 | 1.11 | (-1511.99,-1507.65) |
| 48 | D-Ribulose 5-phosphate | -1598.33 | 1.19 | (-1600.66,-1596.00) |
| 49 | Maltose | -1577.99 | 1.19 | (-1580.33,-1575.65) |
| 50 | Adenosine | -204.39 | 0.41 | (-205.20,-203.57) |
| 51 | D-Arabinose | -741.66 | 0.95 | (-743.51,-739.80) |
| 52 | Adenylyl sulfate | -1525.76 | 1.10 | (-1527.92,-1523.61) |
| 53 | Acetyl phosphate | -1209.83 | 1.06 | (-1211.90,-1207.77) |
| 54 | 4-Methyl-2-oxopentanoate | -447.09 | 0.66 | (-448.38,-445.80) |
| 55 | 3-Phospho-D-glyceroyl phosphate | -2343.14 | 1.47 | (-2346.02,-2340.25) |
| 56 | Lactose | -1574.39 | 1.28 | (-1576.91,-1571.88) |
| 57 | Isomaltose | -1584.23 | 1.20 | (-1586.58,-1581.88) |
| 58 | D-Glycerate | -660.55 | 0.76 | (-662.05,-659.06) |
| 59 | Hypoxanthine | 86.27 | 0.32 | (85.65,86.89) |
| 60 | D-Mannose 6-phosphate | -1761.68 | 1.40 | (-1764.43,-1758.93) |
| 61 | HCO3 | -575.37 | 0.64 | (-576.63,-574.12) |
| 62 | Inosine | -414.49 | 0.60 | (-415.68,-413.31) |
| 63 | D-Ribulose | -735.81 | 0.90 | (-737.58,-734.04) |
| 64 | D-Xylulose | -745.03 | 0.89 | (-746.77,-743.29) |
| 65 | Isocitrate | -1162.28 | 0.94 | (-1164.12,-1160.44) |
| 66 | D-Fructose | -2608.07 | 1.70 | (-2611.41,-2604.74) |
| 67 | cis-Aconitate | -923.66 | 1.03 | (-925.67,-921.65) |
| 68 | Indole | 223.23 | 0.42 | (222.40,224.06) |
| 69 | D-Lyxose | -749.61 | 0.81 | (-751.20,-748.02) |
| 70 | 3',5'-Cyclic AMP | -795.16 | 0.86 | (-796.85,-793.47) |
| 71 | D-Glyceraldehyde | -440.67 | 0.64 | (-441.93,-439.41) |
| 72 | alpha-D-Ribose 1-phosphate | -1583.12 | 1.05 | (-1585.17,-1581.06) |
| 73 | 2-Phospho-D-glycerate | -1502.38 | 1.14 | (-1504.61,-1500.15) |
| 74 | D-Mannose 1-phosphate | -1755.60 | 1.20 | (-1757.95,-1753.26) |
| 75 | 2-Deoxy-D-ribose 5-phosphate | -1451.04 | 1.15 | (-1453.30,-1448.79) |
| 76 | O-Phospho-L-serine | -1364.64 | 1.23 | (-1367.05,-1362.24) |
| 77 | alpha,alpha-Trehalose | -1580.20 | 1.24 | (-1582.64,-1577.76) |
| 78 | D-Arabinose 5-phosphate | -1599.28 | 1.01 | (-1601.27,-1597.29) |
| 79 | D-Galactose 6-phosphate | -1756.01 | 1.35 | (-1758.66,-1753.36) |
| 80 | 4-Hydroxy-2-oxoglutarate | -954.13 | 0.94 | (-955.98,-952.28) |
| 81 | 2-Methylmaleate | -603.63 | 0.76 | (-605.11,-602.15) |
| 82 | (R)-2-Methylmalate | -842.74 | 0.84 | (-844.39,-841.09) |
| 83 | 3-Phosphonooxypyruvate | -1447.42 | 1.15 | (-1449.67,-1445.18) |
| 84 | D-Galactose 1-phosphate | -1757.37 | 1.36 | (-1760.04,-1754.69) |

#### 10.2 Group contribution estimates

The modified group contribution outlined in the main paper (Section 3.2) is used here. In addition to the group contributions, reconciled values are also obtained for the reaction and formation Gibbs free energies.

##### 10.2.1 Confidence intervals of Group Gibbs energy estimates using Group contribution method

| Group Contributions | | | | | |
|---|---|---|---|---|---|
| S.No | Group Name | [H Z Mg] | mean | SE | CI |
| 1 | -SO3 | [0 -1 0] | -649.97 | 0.03 | (-650.71,-649.24) |
| 2 | -CO-OPO3 | [0 -2 0] | -983.75 | 0.02 | (-984.18,-983.32) |
| 3 | -OPO3- | [0 -1 0] | -649.97 | 0.03 | (-650.71,-649.24) |
| 4 | -OPO2- | [0 -1 0] | -292.68 | 0.006 | (-292.81,-292.55) |
| 5 | -OPO3 | [0 -2 0] | -826.82 | 0.01 | (-827.17,-826.46) |
| 6 | -OPO3 | [1 -1 0] | -1422.32 | 0.02 | (-1422.87,-1421.76) |
| 7 | -OPO2-OPO2- | [0 -2 0] | -1129.63 | 0.02 | (-1130.08,-1129.19) |
| 8 | ring -OPO3- | [0 -1 0] | -179.85 | 0.007 | (-179.98,-179.71) |
| 9 | ring =N< | [0 0 0] | -10.05 | 0.015 | (-10.35,-9.75) |
| 10 | ring -N= | [0 0 0] | 117.03 | 0.009 | (116.85,117.21) |
| 11 | N-CO- | [0 0 0] | 3.32 | 0.012 | (3.08,3.56) |
| 12 | -N | [3 1 0] | 165.31 | 0.02 | (164.89,165.74) |
| 13 | -N | [2 0 0] | 3.32 | 0.01 | (3.08,3.56) |
| 14 | ring >C-N | [2 0 0] | 176.38 | 0.01 | (176.02,176.74) |
| 15 | ring >C-O | [1 0 0] | -138.57 | 0.01 | (-138.86,-138.28) |
| 16 | ring >C-O | [2 0 0] | -153.22 | 0.009 | (-153.41,-153.04) |
| 17 | ring >C(-O)- | [1 0 0] | -331.47 | 0.015 | (-331.77,-331.18) |
| 18 | -COO | [0 -1 0] | -150.11 | 0.013 | (-150.36,-149.86) |
| 19 | >C=O | [0 0 0] | -104.28 | 0.02 | (-104.69,-103.87) |
| 20 | -C=O | [1 0 0] | 77.59 | 0.016 | (77.27,77.91) |
| 21 | ring -O- | [0 0 0] | 249.29 | 0.037 | (248.55,250.03) |
| 22 | -C-O | [3 0 0] | 28.24 | 0.01 | (28.03,28.45) |
| 23 | -C(-O)- | [2 0 0] | -152.87 | 0.0048 | (-152.97,-152.78) |
| 24 | >C(-O)- | [1 0 0] | -328.71 | 0.013 | (-328.96,-328.45) |
| 25 | -O- | [0 0 0] | -112.83 | 0.021 | (-113.25,-112.42) |
| 26 | 2-ring -C< | [1 0 0] | -359.70 | 0.01 | (-359.97,-359.43) |
| 27 | 2-ring =C< | [0 0 0] | -12.88 | 0.0074 | (-13.03,-12.74) |
| 28 | ring =C- | [1 0 0] | 87.83 | 0.007 | (87.69,87.96) |
| 29 | ring =C< | [0 0 0] | -101.32 | 0.02 | (-101.75,-100.89) |
| 30 | =C- | [1 0 0] | 45.85 | 0.013 | (45.60,46.10) |
| 31 | =C | [2 0 0] | 204.77 | 0.02 | (204.37,205.17) |
| 32 | =C< | [0 0 0] | -127.10 | 0.016 | (-127.40,-126.79) |
| 33 | ring >C< | [0 0 0] | -341.76 | 0.019 | (-342.14,-341.38) |
| 34 | ring -C< | [1 0 0] | -174.05 | 0.01 | (-174.26,-173.84) |
| 35 | -C< | [1 0 0] | -169.44 | 0.01 | (-169.65,-169.23) |
| 36 | ring -C- | [2 0 0] | 6.06 | 0.012 | (5.82,6.30) |
| 37 | -C- | [2 0 0] | 3.67 | 0.01 | (3.46,3.88) |
| 38 | -C | [3 0 0] | 179.10 | 0.017 | (178.76,179.44) |
| 39 | Origin | [0 0 0] | -393.45 | 0.016 | (-393.76,-393.14) |

##### 10.2.2 Reactions

| Confidence intervals of reaction Gibbs energy estimates using Group contribution method | | | | |
|---|---|---|---|---|
| S.No | Reaction | Mean | SD | CI |
| 1 | isocitrate(aq) = citrate(aq) | -5.24 | 0.07 | (-5.37,-5.11) |

| | | | | |
|---|---|---|---|---|
| 2 | ATP(aq) + D-glucose(aq) = ADP(aq) + D-glucose 6-phosphate(aq) | -10.61 | 0.06 | (-10.72,-10.50) |
| 3 | AMP(aq) + H2O(l) = adenosine(aq) + orthophosphate(aq) | -14.39 | 0.07 | (-14.52,-14.25) |
| 4 | pyrophosphate(aq) + L-serine(aq) = orthophosphate(aq) + O-phospho-L-serine(aq) | 29.01 | 0.10 | (28.82,29.21) |
| 5 | ATP(aq) + D-fructose 6-phosphate(aq) = ADP(aq) + D-fructose 1,6-bisphosphate(aq) | -7.45 | 0.06 | (-7.56,-7.33) |
| 6 | D-fructose 1,6-bisphosphate(aq) = 2 glycerone phosphate(aq) | 10.28 | 0.08 | (10.13,10.43) |
| 7 | D-glucose 6-phosphate(aq) = D-fructose 6-phosphate(aq) | 6.82 | 0.05 | (6.73,6.92) |
| 8 | 2 ADP(aq) = AMP(aq) + ATP(aq) | 37.42 | 0.10 | (37.22,37.63) |
| 9 | inosine(aq) + orthophosphate(aq) = hypoxanthine(aq) + D-ribose 1-phosphate(aq) | 9.13 | 0.07 | (8.99,9.26) |
| 10 | D-fructose 1,6-bisphosphate(aq) + H2O(l) = D-fructose 6-phosphate(aq) + orthophosphate(aq) | -13.86 | 0.05 | (-13.96,-13.76) |
| 11 | ATP(aq) + 3-phospho-D-glycerate(aq) = ADP(aq) + 3-phospho-D-glyceroyl phosphate(aq) | 10.64 | 0.08 | (10.47,10.80) |
| 12 | D-glyceraldehyde 3-phosphate(aq) = glycerone phosphate(aq) | -5.62 | 0.06 | (-5.75,-5.50) |
| 13 | ATP(aq) + pyruvate(aq) + orthophosphate(aq) = AMP(aq) + phosphoenolpyruvate(aq) + pyrophosphate(aq) | 15.39 | 0.08 | (15.23,15.56) |
| 14 | D-xylose(aq) = D-xylulose(aq) | 3.87 | 0.06 | (3.76,3.98) |
| 15 | ATP(aq) + L-glutamate(aq) + ammonia(aq) = ADP(aq) + phosphate(aq) + L-glutamine(aq) | -15.58 | 0.08 | (-15.74,-15.42) |
| 16 | D-mannose 6-phosphate(aq) = D-fructose 6-phosphate(aq) | 2.89 | 0.04 | (2.82,2.97) |
| 17 | ATP(aq) = adenosine 3':5'-(cyclic)phosphate(aq) + diphosphate(aq) | 7.34 | 0.08 | (7.18,7.50) |
| 18 | sucrose(aq) + orthophosphate(aq) =-D-glucose 1-phosphate(aq) + D-fructose(aq) | -10.11 | 0.10 | (-10.30,-9.91) |
| 19 | L-alanine(aq) + 2-oxoglutarate(aq) = pyruvate(aq) + L-glutamate(aq) | 2.49 | 0.04 | (2.41,2.58) |
| 20 | diphosphate(aq) + oxaloacetate(aq) + H2O(l) = phosphate(aq) + phosphoenolpyruvate(aq) + carbon dioxide(aq) | 39.19 | 0.10 | (38.99,39.40) |
| 21 | D-mannose(aq) = D-fructose(aq) | -1.91 | 0.03 | (-1.96,-1.85) |
| 22 | isocitrate(aq) = succinate(aq) + glyoxylate(aq) | 7.81 | 0.08 | (7.66,7.96) |
| 23 | D-galactose 1-phosphate(aq) =-D-glucose 1-phosphate(aq) | -1.34 | 0.04 | (-1.41,-1.27) |
| 24 | fumarate(aq) + H2O(l) = (S)-malate(aq) | -15.11 | 0.12 | (-15.34,-14.88) |
| 25 | D-glucose(aq) = D-fructose(aq) | 0.26 | 0.02 | (0.23,0.29) |
| 26 | 2-phospho-D-glycerate(aq) = 3-phospho-D-glycerate(aq) | -7.84 | 0.08 | (-7.99,-7.69) |
| 27 | D-fructose 6-phosphate(aq) + H2O(l) = D-fructose(aq) + orthophosphate(aq) | -17.26 | 0.05 | (-17.36,-17.16) |
| 28 | 3-phosphonooxypyruvate(aq) + L-glutamate(aq) = 2-oxoglutarate(aq) + O-phospho-L-serine(aq) | -9.41 | 0.06 | (-9.53,-9.29) |

| | | | | |
|---|---|---|---|---|
| 29 | adenosine(aq) + H2O(l) = adenine(aq) + D-ribose(aq) | -15.19 | 0.08 | (-15.35,-15.03) |
| 30 | L-aspartate(aq) + 2-oxoglutarate(aq) = oxaloacetate(aq) + L-glutamate(aq) | -0.64 | 0.03 | (-0.71,-0.58) |
| 31 | -D-glucose 1-phosphate(aq) =-D-glucose 6-phosphate(aq) | -5.67 | 0.05 | (-5.76,-5.58) |
| 32 | ATP(aq) + D-ribose 5-phosphate(aq) = AMP(aq) + 5-phospho–D-ribose 1-diphosphate(aq) | -16.79 | 0.09 | (-16.98,-16.61) |
| 33 | L-aspartate(aq) = fumarate(aq) + ammonia(aq) | 15.21 | 0.11 | (15.00,15.43) |
| 34 | AMP(aq) + H2O(l) = adenine(aq) + D-ribose 5-phosphate(aq) | -20.32 | 0.08 | (-20.48,-20.17) |
| 35 | L-leucine(aq) + 2-oxoglutarate(aq) = 2-oxoisocaproate(aq) + L-glutamate(aq) | 0.63 | 0.03 | (0.57,0.68) |
| 36 | (R)-2-methylmalate(aq) = 2-methylmaleate(aq) + H2O(l) | 14.62 | 0.09 | (14.45,14.80) |
| 37 | D-fructose 1,6-bisphosphate(aq) = glycerone phosphate(aq) + D-glyceraldehyde 3-phosphate(aq) | 15.90 | 0.09 | (15.72,16.08) |
| 38 | 4-hydroxy-2-oxoglutarate(aq) = pyruvate(aq) + glyoxylate(aq) | 14.06 | 0.10 | (13.86,14.26) |
| 39 | lactose(aq) + H2O(l) = D-galactose(aq) + D-glucose(aq) | -11.16 | 0.08 | (-11.32,-11.01) |
| 40 | D-ribose 1-phosphate(aq) = D-ribose 5-phosphate(aq) | -11.70 | 0.06 | (-11.81,-11.58) |
| 41 | citrate(aq) = cis-aconitate(aq) + H2O(l) | 18.26 | 0.08 | (18.10,18.42) |
| 42 | D-ribose 5-phosphate(aq) = D-ribulose 5-phosphate(aq) | 4.79 | 0.06 | (4.67,4.90) |
| 43 | D-glucose 6-phosphate(aq) + H2O(l) = D-glucose(aq) + orthophosphate(aq) | -10.70 | 0.05 | (-10.79,-10.60) |
| 44 | citrate(aq) = acetate(aq) + oxaloacetate(aq) | 3.76 | 0.05 | (3.66,3.87) |
| 45 | alpha,alpha-Trehalose + H2O(l) = 2 D-glucose(aq) | -11.78 | 0.07 | (-11.91,-11.65) |
| 46 | isocitrate(aq) = cis-aconitate(aq) + H2O(l) | 13.02 | 0.08 | (12.87,13.17) |
| 47 | ATP(aq) + pyruvate(aq) + carbon dioxide(aq) = ADP(aq) + phosphate(aq) + oxaloacetate(aq) | -2.38 | 0.05 | (-2.47,-2.29) |
| 48 | 2-phospho-D-glycerate(aq) = phosphoenolpyruvate(aq) + H2O(l) | -8.71 | 0.08 | (-8.86,-8.55) |
| 49 | ATP(aq) + sulfate(aq) = adenosine 5'-phosphosulfate(aq) + pyrophosphate(aq) | 38.72 | 0.16 | (38.40,39.03) |
| 50 | adenine(aq) + 5-phospho–D-ribose 1-diphosphate(aq) = AMP(aq) + pyrophosphate(aq) | 13.16 | 0.07 | (13.02,13.30) |
| 51 | isomaltose(aq) + H2O(l) = 2 D-glucose(aq) | -8.48 | 0.07 | (-8.62,-8.33) |
| 52 | AMP(aq) + pyrophosphate(aq) = adenine(aq) + 5-phospho–D-ribose 1-diphosphate(aq) | -13.16 | 0.07 | (-13.30,-13.02) |
| 53 | 2-deoxy-D-ribose 5-phosphate(aq) = D-glyceraldehyde 3-phosphate(aq) + acetaldehyde(aq) | 16.31 | 0.13 | (16.06,16.56) |
| 54 | L-glutamine(aq) + H2O(l) = L-glutamate(aq) + ammonia(aq) | -5.73 | 0.07 | (-5.87,-5.58) |
| 55 | ATP(aq) + pyruvate(aq) = ADP(aq) + phosphoenolpyruvate(aq) | 18.04 | 0.08 | (17.88,18.20) |
| 56 | adenosine(aq) + orthophosphate(aq) = adenine(aq) + D-ribose 1-phosphate(aq) | 5.76 | 0.07 | (5.62,5.89) |

| | | | | |
|---|---|---|---|---|
| 57 | D-arabinose 5-phosphate(aq) = D-ribulose 5-phosphate(aq) | 1.68 | 0.04 | (1.60,1.76) |
| 58 | O-phospho-L-serine(aq) + 2-oxoglutarate(aq) = 3-phosphonooxypyruvate(aq) + L-glutamate(aq) | 9.41 | 0.06 | (9.29,9.53) |
| 59 | D-lyxose(aq) = D-xylulose(aq) | 3.03 | 0.05 | (2.93,3.13) |
| 60 | L-tryptophan(aq) + H2O(l) = indole(aq) + pyruvate(aq) + ammonia(aq) | 22.67 | 0.16 | (22.35,22.98) |
| 61 | (S)-lactate(aq) + oxaloacetate(aq) = (S)-malate(aq) + pyruvate(aq) | -0.40 | 0.02 | (-0.45,-0.36) |
| 62 | pyruvate(aq) + orthophosphate(aq) = acetyl phosphate(aq) + formate(aq) | -6.86 | 0.08 | (-7.02,-6.70) |
| 63 | D-mannose 6-phosphate(aq) + H2O(l) = D-mannose(aq) + orthophosphate(aq) | -12.46 | 0.05 | (-12.57,-12.36) |
| 64 | D-mannose 1-phosphate(aq) = D-mannose 6-phosphate(aq) | -3.79 | 0.05 | (-3.89,-3.69) |
| 65 | ATP(aq) + acetate(aq) = ADP(aq) + acetyl phosphate(aq) | 9.94 | 0.08 | (9.77,10.10) |
| 66 | (R)-3-phosphoglycerate(aq) + H2O(l) = (R)-glycerate(aq) + orthophosphate(aq) | -14.77 | 0.07 | (-14.91,-14.63) |
| 67 | trehalose(aq) + orthophosphate(aq) = D-glucose(aq) + D-glucose 1-phosphate(aq) | 4.59 | 0.05 | (4.48,4.69) |
| 68 | glycine(aq) + formaldehyde(aq) = L-serine(aq) | -16.98 | 0.11 | (-17.20,-16.77) |
| 69 | D-ribose(aq) = D-ribulose(aq) | 1.11 | 0.03 | (1.05,1.17) |
| 70 | adenosine(aq) + H2O(l) = inosine(aq) + ammonia(aq) | -62.37 | 0.20 | (-62.75,-61.98) |
| 71 | pyrophosphate(aq) + H2O(l) = 2 orthophosphate(aq) | 18.77 | 0.07 | (18.64,18.90) |
| 72 | D-fructose(aq) + D-glyceraldehyde-3-phosphate(aq) = D-fructose 6-phosphate(aq) + D-glyceraldehyde(aq) | 3.36 | 0.05 | (3.27,3.45) |
| 73 | ATP(aq) + sulfate(aq) + H2O(l) = 2 orthophosphate(aq) + adenosine 5'-phosphosulfate(aq) | 57.49 | 0.16 | (57.18,57.79) |
| 74 | ATP(aq) + H2O(l) = ADP(aq) + orthophosphate(aq) | -21.31 | 0.05 | (-21.41,-21.21) |
| 75 | ATP(aq) + pyruvate(aq) + H2O(l) = AMP(aq) + phosphoenolpyruvate(aq) + orthophosphate(aq) | 34.16 | 0.08 | (33.99,34.33) |
| 76 | maltose(aq) + orthophosphate(aq) = D-glucose(aq) + D-glucose 1-phosphate(aq) | 2.11 | 0.04 | (2.03,2.19) |
| 77 | maltose(aq) + H2O(l) = 2 D-glucose(aq) | -14.26 | 0.06 | (-14.37,-14.14) |
| 78 | pyrophosphate(aq) + D-fructose 6-phosphate(aq) = orthophosphate(aq) + D-fructose 1,6-bisphosphate(aq) | 32.63 | 0.07 | (32.48,32.77) |
| 79 | D-galactose 6-phosphate(aq) + H2O(l) = D-galactose(aq) + orthophosphate(aq) | -11.55 | 0.07 | (-11.69,-11.42) |
| 80 | L-O-phosphoserine(aq) + H2O(l) = L-serine(aq) + orthophosphate(aq) | -10.25 | 0.07 | (-10.38,-10.11) |
| 81 | D-arabinose(aq) = D-ribulose(aq) | 5.34 | 0.04 | (5.26,5.43) |
| 82 | glycine(aq) + acetaldehyde(aq) = L-threonine(aq) | -11.74 | 0.08 | (-11.90,-11.58) |
| 83 | inosine(aq) + H2O(l) = hypoxanthine(aq) + D-ribose(aq) | -11.83 | 0.07 | (-11.96,-11.69) |

| | | | | |
|---|---|---|---|---|
| 84 | D-arabinose(aq) = D-ribose(aq) | 4.23 | 0.05 | (4.14,4.32) |
| 85 | D-glucose(aq) = D-mannose(aq) | 2.17 | 0.03 | (2.11,2.22) |
| 86 | ATP(aq) + D-galactose(aq) = ADP(aq) + D-galactose 1-phosphate(aq) | -6.07 | 0.06 | (-6.19,-5.94) |
| 87 | glycine(aq) + oxaloacetate(aq) = glyoxylate(aq) + L-aspartate(aq) | 6.56 | 0.06 | (6.45,6.68) |

##### 10.2.3 Compounds

| Confidence intervals of estimates formation Gibbs energies using group contribution method | | | | |
|---|---|---|---|---|
| S.No | Compound | Mean | SE | CI |
| 1 | H2O | -227.35 | 0.17 | (-227.69,-227.02) |
| 2 | ATP | -2781.00 | 0.55 | (-2782.08,-2779.93) |
| 3 | ADP | -1937.86 | 0.41 | (-1938.66,-1937.06) |
| 4 | Orthophosphate | -1091.81 | 0.23 | (-1092.26,-1091.35) |
| 5 | Diphosphate | -1975.03 | 0.36 | (-1975.74,-1974.31) |
| 6 | NH3 | -76.79 | 0.21 | (-77.21,-76.37) |
| 7 | AMP | -1057.29 | 0.29 | (-1057.85,-1056.73) |
| 8 | Pyruvate | -467.60 | 0.12 | (-467.83,-467.37) |
| 9 | L-Glutamate | -690.00 | 0.23 | (-690.45,-689.55) |
| 10 | 2-Oxoglutarate | -791.17 | 0.25 | (-791.67,-790.67) |
| 11 | D-Glucose | -903.55 | 0.20 | (-903.94,-903.15) |
| 12 | Acetate | -366.42 | 0.18 | (-366.77,-366.06) |
| 13 | Oxaloacetate | -795.78 | 0.27 | (-796.31,-795.26) |
| 14 | Glycine | -372.34 | 0.20 | (-372.74,-371.95) |
| 15 | L-Alanine | -368.92 | 0.19 | (-369.29,-368.55) |
| 16 | Succinate | -685.32 | 0.21 | (-685.74,-684.89) |
| 17 | Glyoxylate | -467.60 | 0.17 | (-467.94,-467.26) |
| 18 | L-Aspartate | -693.97 | 0.23 | (-694.42,-693.52) |
| 19 | Formate | -366.64 | 0.22 | (-367.06,-366.22) |
| 20 | Sulfate | -761.74 | 0.76 | (-763.23,-760.26) |
| 21 | L-Glutamine | -533.71 | 0.36 | (-534.41,-533.01) |
| 22 | L-Serine | -519.81 | 0.19 | (-520.20,-519.43) |
| 23 | Formaldehyde | -130.49 | 0.13 | (-130.74,-130.23) |
| 24 | Phosphoenolpyruvate | -1292.70 | 0.20 | (-1293.10,-1292.30) |
| 25 | L-Tryptophan | -114.94 | 0.27 | (-115.48,-114.41) |
| 26 | Acetaldehyde | -136.70 | 0.12 | (-136.93,-136.46) |
| 27 | D-Fructose 6-phosphate | -1750.48 | 0.22 | (-1750.92,-1750.05) |
| 28 | Sucrose | -1553.01 | 0.39 | (-1553.76,-1552.25) |
| 29 | D-Glucose 6-phosphate | -1757.30 | 0.22 | (-1757.73,-1756.87) |
| 30 | D-Fructose | -903.29 | 0.20 | (-903.68,-902.90) |
| 31 | D-Glucose | -1751.63 | 0.22 | (-1752.07,-1751.20) |
| 32 | Glycerone phosphate | -1295.40 | 0.17 | (-1295.73,-1295.07) |
| 33 | D-Ribose 5-phosphate | -1602.54 | 0.20 | (-1602.94,-1602.15) |
| 34 | D-Glyceraldehyde 3-phosphate | -1289.77 | 0.17 | (-1290.12,-1289.43) |
| 35 | 5-Phospho-alpha-D-ribose 1-diphosphate | -3343.05 | 0.48 | (-3344.00,-3342.11) |
| 36 | D-Ribose | -747.35 | 0.19 | (-747.72,-746.98) |
| 37 | Fumarate | -601.96 | 0.25 | (-602.45,-601.47) |

| | | | | |
|---|---|---|---|---|
| 38 | L-Leucine | -354.91 | 0.36 | (-355.62,-354.19) |
| 39 | D-Galactose | -901.08 | 0.20 | (-901.47,-900.69) |
| 40 | Adenine | 297.58 | 0.28 | (297.04,298.12) |
| 41 | (S)-Malate | -844.43 | 0.20 | (-844.81,-844.04) |
| 42 | Citrate | -1165.96 | 0.29 | (-1166.53,-1165.40) |
| 43 | D-Mannose | -901.38 | 0.20 | (-901.77,-900.99) |
| 44 | D-Xylose | -751.47 | 0.19 | (-751.85,-751.09) |
| 45 | (S)-Lactate | -515.84 | 0.19 | (-516.22,-515.46) |
| 46 | L-Threonine | -520.78 | 0.19 | (-521.15,-520.41) |
| 47 | 3-Phospho-D-glycerate | -1519.19 | 0.23 | (-1519.65,-1518.73) |
| 48 | D-Ribulose 5-phosphate | -1597.76 | 0.21 | (-1598.18,-1597.34) |
| 49 | Maltose | -1565.49 | 0.40 | (-1566.27,-1564.70) |
| 50 | Adenosine | -207.22 | 0.25 | (-207.72,-206.73) |
| 51 | D-Arabinose | -751.58 | 0.19 | (-751.96,-751.20) |
| 52 | Adenylyl sulfate | -1529.00 | 0.82 | (-1530.60,-1527.40) |
| 53 | Acetyl phosphate | -1199.63 | 0.27 | (-1200.16,-1199.09) |
| 54 | 4-Methyl-2-oxopentanoate | -455.45 | 0.31 | (-456.05,-454.85) |
| 55 | 3-Phospho-D-glyceroyl phosphate | -2351.70 | 0.38 | (-2352.44,-2350.95) |
| 56 | Lactose | -1566.11 | 0.40 | (-1566.89,-1565.34) |
| 57 | Isomaltose | -1571.26 | 0.39 | (-1572.03,-1570.50) |
| 58 | D-Glycerate | -669.50 | 0.18 | (-669.85,-669.16) |
| 59 | Hypoxanthine | 88.02 | 0.27 | (87.49,88.55) |
| 60 | D-Mannose 6-phosphate | -1753.37 | 0.22 | (-1753.80,-1752.95) |
| 61 | HCO3 | -574.46 | 0.24 | (-574.92,-574.00) |
| 62 | Inosine | -420.15 | 0.28 | (-420.70,-419.60) |
| 63 | D-Ribulose | -746.24 | 0.19 | (-746.61,-745.86) |
| 64 | D-Xylulose | -747.60 | 0.19 | (-747.98,-747.22) |
| 65 | Isocitrate | -1160.72 | 0.29 | (-1161.29,-1160.16) |
| 66 | D-Fructose | -2601.08 | 0.34 | (-2601.73,-2600.42) |
| 67 | cis-Aconitate | -920.35 | 0.33 | (-921.00,-919.71) |
| 68 | Indole | 224.76 | 0.33 | (224.11,225.42) |
| 69 | D-Lyxose | -750.63 | 0.20 | (-751.02,-750.24) |
| 70 | 3',5'-Cyclic AMP | -798.64 | 0.39 | (-799.41,-797.86) |
| 71 | D-Glyceraldehyde | -439.22 | 0.13 | (-439.47,-438.97) |
| 72 | alpha-D-Ribose 1-phospate | -1590.85 | 0.21 | (-1591.25,-1590.45) |
| 73 | 2-Phospho-D-glycerate | -1511.35 | 0.22 | (-1511.79,-1510.91) |
| 74 | D-Mannose 1-phosphate | -1749.59 | 0.22 | (-1750.01,-1749.16) |
| 75 | 2-Deoxy-D-ribose 5-phosphate | -1442.78 | 0.25 | (-1443.27,-1442.29) |
| 76 | O-Phospho-L-serine | -1374.02 | 0.24 | (-1374.49,-1373.55) |
| 77 | alpha,alpha-Trehalose | -1567.96 | 0.39 | (-1568.72,-1567.20) |
| 78 | D-Arabinose 5-phosphate | -1599.44 | 0.22 | (-1599.87,-1599.01) |
| 79 | D-Galactose 6-phosphate | -1753.98 | 0.22 | (-1754.42,-1753.55) |
| 80 | 4-Hydroxy-2-oxoglutarate | -949.26 | 0.26 | (-949.76,-948.75) |
| 81 | 2-Methylmaleate | -596.81 | 0.24 | (-597.28,-596.34) |
| 82 | (R)-2-Methylmalate | -838.78 | 0.22 | (-839.21,-838.35) |
| 83 | 3-Phosphonooxypyruvate | -1465.79 | 0.22 | (-1466.22,-1465.35) |
| 84 | D-Galactose 1-phosphate | -1750.29 | 0.22 | (-1750.72,-1749.87) |

[2] Alberty. *Thermodynamics of biochemical reactions.* John Wiley & Sons, 2005.

[3] CM Crowe, YA Campos, and A Hrymak. Reconciliation of process flow rates by matrix projection. part i: Linear case. *AIChE Journal*, 29(6):881–888, 1983.

[4] Narasimhan et al. *Data reconciliation and gross error detection: An intelligent use of process data.* Gulf Professional Publishing, 1999.

[5] Noor et al. An integrated open framework for thermodynamics of reactions that combines accuracy and coverage. *Bioinformatics*, 28(15):2037–2044, 2012.

[6] Noor et al. Consistent estimation of gibbs energy using component contributions. *PLoS computational biology*, 9(7):e1003098, 2013.
